# Protein-Induced Membrane Strain Drives Supercomplex Formation

**DOI:** 10.1101/2024.07.13.602826

**Authors:** Maximilian C. Pöverlein, Alexander Jussupow, Hyunho Kim, Ville R. I. Kaila

## Abstract

Mitochondrial membranes harbor the electron transport chain (ETC) that powers oxidative phosphorylation (OXPHOS) and drives the synthesis of ATP. Yet, under physiological conditions, the OXPHOS proteins operate as higher-order supercomplex (SC) assemblies, although their functional role remains poorly understood and much debated. By combining large-scale atomistic and coarse-grained molecular simulations with analysis of cryo-electron microscopic data and statistical as well as kinetic models, we show here that the formation of the mammalian I/III_2_ supercomplex reduces the molecular strain of inner mitochondrial membranes by altering the local membrane thickness and leading to an accumulation of both cardiolipin and quinone around specific regions of the SC. We find that the SC assembly also affects the global motion of the individual ETC proteins with possible functional consequences. On a general level, our findings suggest that molecular crowding and strain effects provide a thermodynamic driving force for the SC formation, with a possible flux enhancement in crowded biological membranes under constrained respiratory conditions.

**Significance Statement:** The membrane-bound proteins of respiratory chains power oxidative phosphorylation (OXPHOS) and drive the synthesis of ATP. However, recent biochemical and structural data show that the OXPHOS proteins operate as higher-order supercomplex assemblies for reasons that remain elusive and much debated. Here we show that the mammalian respiratory supercomplexes reduce the molecular strain of inner mitochondrial membranes and enhance the allosteric crosstalk by altering the protein dynamics with important biochemical and physiological implications.

## Introduction

Biological electron transport chains (ETC) comprise a series of membrane-bound enzyme complexes (CI-CIV) that transfer electrons towards oxygen and protons across a biological membrane, creating a proton motive force (PMF) that powers the synthesis of ATP and active transport (1, 2). Biological membranes are often envisaged in the light of the classical fluid mosaic model (3), where the membrane proteins and lipids independently diffuse as a 2-dimensional solution. However, the biological membranes are highly crowded (4-6), particularly the inner mitochondrial membrane (IMM), with a protein content of around 45% (7). In this regard, the majority of the respiratory complexes do not diffuse independently within the membrane but operate rather as larger supercomplex (SC) assemblies, as first revealed by blue native gels (8) and confirmed by structural analyses (9-11). Moreover, several lipid molecules, particularly cardiolipin, a central anionic lipid of the IMM, is often tightly bound to the SCs (12-15). Yet, despite these structural insights, the functional role of the SC assemblies and the physical principles leading to their formation remain elusive and much debated.

SCs have been suggested to provide kinetic advantages (12, 16), favor substrate channeling ((10, 17), but *cf*. (18)), decrease the formation of reactive oxygen species (ROS) (19, 20), stabilize individual proteins (21-24), and/or reduce non-specific protein-protein interactions (18). It has also been recognized that the high protein density in membranes may play an important role in the SC formation (18), and influence, *e*.*g*., the lipid curvature (7, 25). On a physiological level, decreased SC formation has been linked to various patho-physiological conditions such as diabetes (26), heart failure (27), and apoptosis (28), although recent experiments (29) on mice unable to form SC showed no significant differences relative to WT mice under the studied conditions.

In recent years, cryo-electron microscopic (cryo-EM) and cryo-electron tomographic (cryo-TEM) studies revealed the structure of several SCs, including the mammalian 1.5 MDa SCI/III_2_ and the 1.7 MDa respirasome (SCI/III_2_/IV) (30-34), as well as various exotic megacomplexes isolated from single-celled eukaryotes (35, 36). SCs are also essential for some bacteria, such as the SCIII_2_/IV_2_ of actinobacteria (15, 37-39) that catalyze quinol oxidation coupled to oxygen reduction in a tight obligate protein assembly. Inhibition of such bacterial SCs provides potential avenues for treating pathogenic infections, such as tuberculosis, with emerging multi-drug resistant strains.

To probe the physical principles leading to the SC formation and its molecular consequences, we study here the structure and dynamics of the mammalian mitochondrial SCI/III_2_ by combining large-scale atomistic and coarse-grained molecular dynamics simulations with analysis of cryo-EM data and mathematical models that provide insight into crowding effects, membrane strain, deformation effects, as well as dynamic protein-lipid interactions influence SC formation.

The studied SCI/III_2_ catalyzes an NADH-driven quinone / cytochrome *c* reduction and offers a unique system to probe how the quinone (Q) substrate is transported within the SC assembly, and how the individual proteins interact within the membrane. In this regard, the quinol (QH_2_) produced by the CI module of the SC is re-oxidized by the CIII module, whilst the proton transport activity (CI: 2H^+^/e^−^; CIII_2_: 1H^+^/e^−^) powers the synthesis of ATP (Fig. 1). Interestingly, prior cryo-EM studies of this SC (40) revealed an asymmetric protein assembly with a trapped quinone in the proximal/distal Q_o_ site, suggesting possible functional consequences. However, the dynamics and interaction of the SC and its substrates within the surrounding membrane environment and the individual active site remain poorly understood.

**Figure 1.**
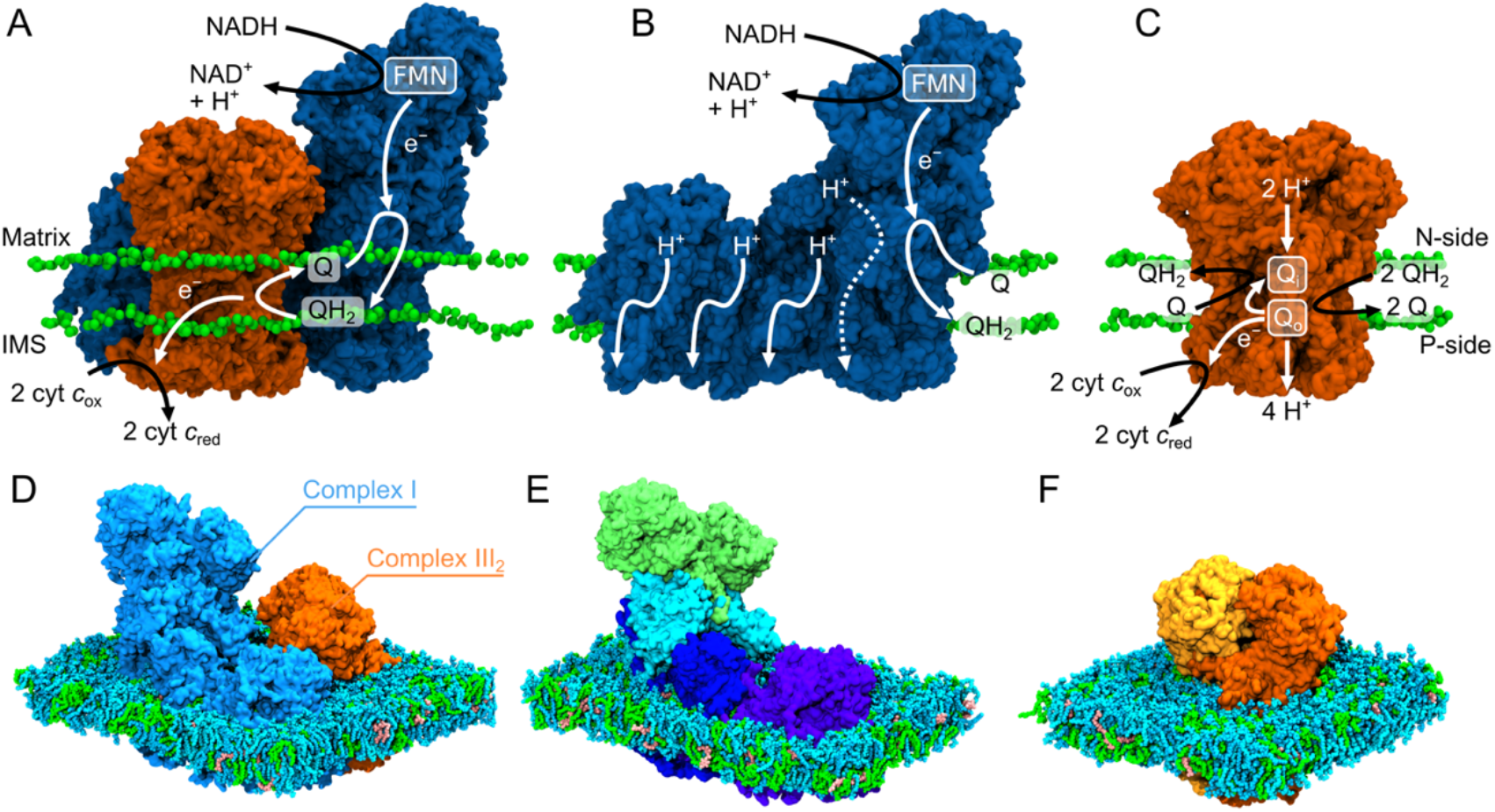
Structure and function of Complex I, Complex III_2_, and the SCI/III_2_. (**A**) The structure of the OXPHOS complexes with lipid headgroups from atomistic MD simulations. (**B, C**) Overview of the structure and function of redox-driven proton pumping in CI and the Q-cycle of CIII_2_. (**D**-**F**) Molecular models of the (**D**) SCI/III_2_, (**E**) CI, and (**F**) CIII_2_ used in the atomistic MD simulations (water and ions were omitted for visual clarity).

## Results

### Protein-membrane interactions modulate the lipid dynamics and membrane strain

To probe how the SC formation affects the dynamics of the OXPHOS proteins, the lipid membrane, and the quinone pool, we studied the ovine SCI/III_2_ by both atomistic (aMD) and coarse-grained molecular dynamics (cgMD) simulations. While the aMD simulations give insight into the microsecond dynamics (11 μs in total, Table S1) of the system with atomic details, our cgMD simulations allowed us to probe the SC dynamics and its interactions in the membrane on much longer timescales (∼0.3 ms in cumulative simulation time, 50-75 μs/simulation) and with larger membrane size at a more approximate level (see *Methods*). The SC models were further compared against simulations of the individual CI and CIII_2_ complexes, as well as simulations of a lipid membrane, with a POPC:POPE:CDL ratio (2:2:1) as in the IMM. In the following, we define *membrane strain* as the local perturbations of the lipid bilayer induced by protein-membrane interactions. These include changes in (i) membrane thickness,(ii) the local membrane composition, (iii) lipid chain configurations, and (iv) local curvature of the membrane plane relative to an undisturbed, protein-free bilayer. Together, these phenomena reflect the thermodynamic effects associated with accommodating large protein complexes within the membrane.

We find that the individual OXPHOS complexes, CI and CIII_2_, induce pronounced membrane strain effects, supported both by our aMD (Fig. S2A) and cgMD simulations with a large surrounding membrane (Fig. 2G). These effects persist for both individual complexes and assembled SC. We observe a local decrease in the membrane thickness at the protein-lipid interface (Fig. 2G, Fig S2A,D,E), likely arising from the thinner hydrophobic belt region of the OXPHOS proteins (*ca*. 30 Å, Fig. S1A) relative to the lipid membrane (40.5 Å, Fig. S1D). We further observe ∼30% accumulation of cardiolipin at the thinner hydrophobic belt regions (Fig. 2H, Fig. S2B,F,G), with an inhomogeneous distribution around the OXPHOS complexes. While specific interactions between CDL and protein residues may contribute to this enrichment (Fig. 2N), CDL also thermodynamically prefers thinner membranes (∼38 Å, Fig. S1D, Fig. S5E). These changes are further reflected in the reduced *end*-to-*end* distance of lipid chains in the local membrane belt (see *Methods*, Fig. S6, cf. also Refs. (41-44)). In addition to the perturbations in the local membrane thickness, the OXPHOS proteins also induce a subtle inward curvature towards the protein-lipid interface (Fig. S5G), which could modulate the accessibility of the Q/QH_2_ substrate into the active sites of CI and CIII_2_ (see below, section *Discussion*). This curvature is accompanied by a distortion of the local membrane plane itself (Fig. 2A-F, Fig. S4A-C, Fig. S8), with perpendicular leaflet displacements reaching up to ∼2 nm relative to the average leaflet plane.

**Figure 2.**
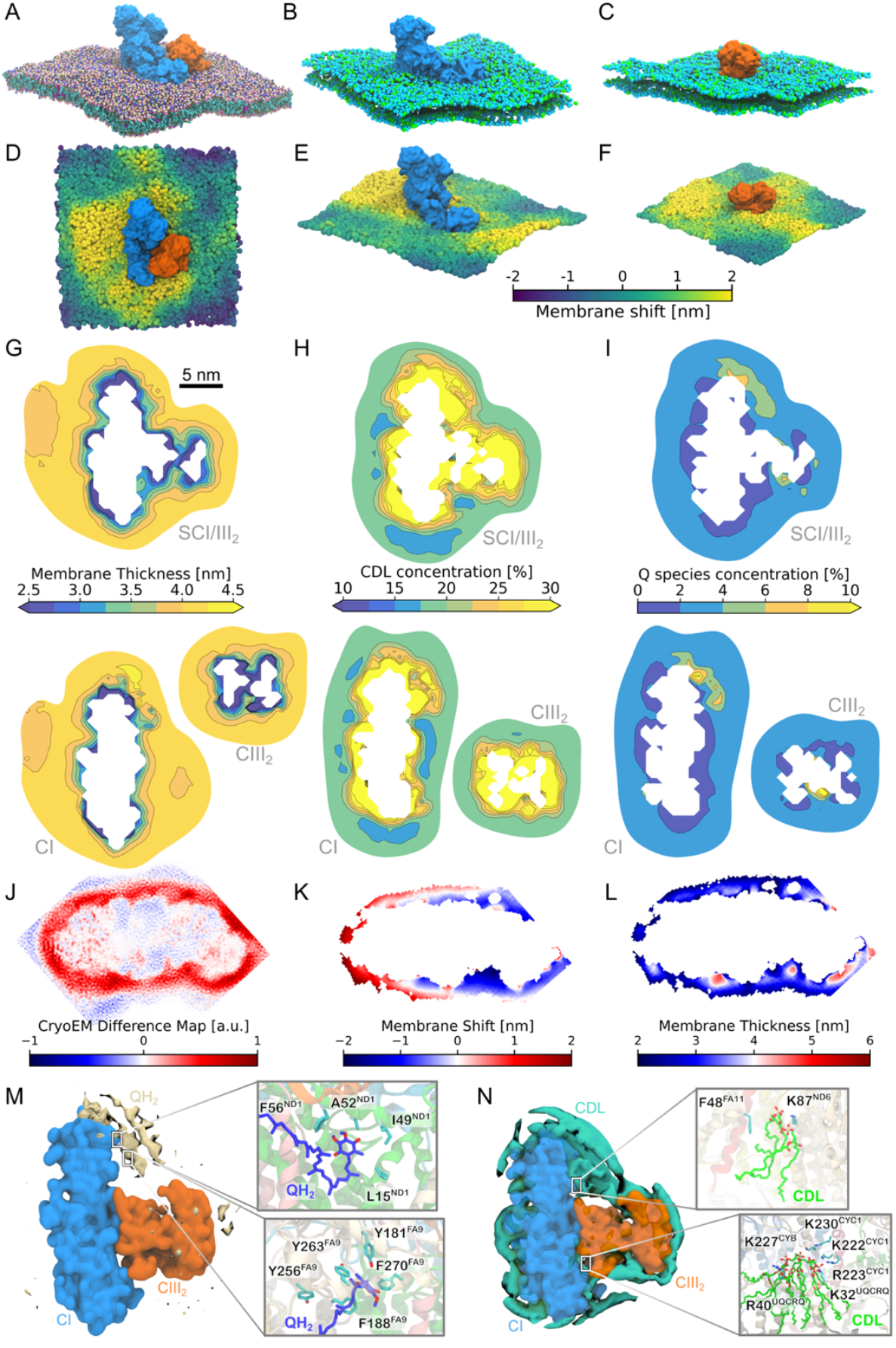
SC formation affects the concentration of lipids and the membrane thickness. (**A-C**) OXPHOS complexes viewed from the N-side of the membrane. (**D-F**) Membrane shift induced by (**D**) the SC, and (**E**,**F**) the isolated CI and CIII_2_. The membrane shift relative to the average membrane plane is colored by the shift in *z*-position viewed from the N-side. (**G**) Local membrane thickness around SC (top), CI and CIII_2_ (bottom). (**H**) Local cardiolipin concentration. (**I**) Local concentration of Q (Q+QH_2_). (**J**) Cryo-EM difference map. (**K**) Membrane shift determined from the cryo-EM map. (**L**) Membrane thickness determined from cryo-EM map. (**M**) Density of Q/QH_2_ (headgroup) from cgMD simulations. *Insets:* Protein-Q interactions from atomistic MD. (**N**) Density of cardiolipin (headgroup) from cgMD simulations. *Insets:* Protein-CDL interactions from atomistic MD.

To quantify the membrane strain effects, we analyzed the cgMD trajectories by projecting the membrane surface onto a 2-dimensional grid and calculating the local membrane height and thickness at each grid point. From these values, we quantified the *local membrane curvature* (Fig. S5H), which measures the energetic cost of bending the membrane from a flat geometry (Δ*G*_curv_) based on the Helfrich model (45, 46). We also computed the energetics associated with changes in the *membrane thickness*, assessed from the deviations from an ideal local membrane in the absence of embedded proteins (Δ*G*_thick_, see *Supporting Information*, for technical details). Our analysis suggests that both contributions are substantially reduced upon formation of the SC, with the curvature penalty decreasing by 79.2 ± 5.2 kcal mol^-1^ (for a membrane area of ca. 1000 nm^2^) and the thickness penalty by 2.8 ± 2.0 kcal mol^-1^ (Table 1). These results indicate a significant thermodynamic advantage for SC formation, as it minimizes lipid deformation and stabilizes the membrane environment surrounding Complex I and III.

**Table 1.** Estimated membrane deformation energies associated with individual complexes and their supercomplex (SC). Energies are reported as mean ± standard deviation from bootstrap analysis. The deformation energies are reported for a membrane patch with an area of *ca*. 1000 nm^2^.

| | $\Delta G_{\text{curv}}$ (kcal mol<sup>-1</sup>) | $\Delta G_{\text{thick}}$ (kcal mol<sup>-1</sup>) |
| --- | --- | --- |
| CI | 123.6 ± 2.8 | 23.9 ± 0.6 |
| CIII<sub>2</sub> | 59.6 ± 4.4 | 28.0 ± 1.4 |
| SC | 103.6 ± 1.6 | 49.1 ± 0.9 |
| **SC – (CI + CIII<sub>2</sub>)** | **-79.2 ± 5.2** | **-2.8 ± 2.0** |

Remarkably, the SC assembly partially releases this strain energy as a result of a smaller solvation area established at the SC-lipid interface (Fig. 2G, Fig. S5, Fig. S8, Table S4). We find that an increase in the lipid tail length decreases the relative stability of the SC (Fig. S5F), further supporting that the hydrophobic mismatch between the OXPHOS proteins and the lipid membrane modulates the strain effect (Fig. S5A-C).

Taken together, the analysis suggests that the OXPHOS complexes affect the mechanical properties of the membranes by inducing a small inwards curvature towards the protein-lipid interface (Fig. S5G,H), resulting in a membrane deformation effect, while the SC formation releases some deformation energy relative to the isolated OXPHOS complexes. The localization of specific lipids around the membrane proteins, as well as local membrane perturbation effects, is also supported by simulations of other membrane proteins (47, 48), suggesting that the effects could arise from general protein-membrane interactions.

Although the current cryo-EM maps do not resolve the molecular details of the surrounding lipid membrane (but see below), we performed a spatial integration of the experimental cryo-EM density map by training a neural network model (see *Methods*) that allowed us to identify and calculate the membrane thickness of the lipid belt around Complex I (PDB ID: 6RFR, EMD-4873) (49). In this regard, we find that the experimental data also shows a statistically significant membrane thinning around CI relative to the SC at the protein-membrane interface (Fig. 2J-L), strongly supporting our simulation results. We note that during the finalization of this work, a membrane distortion effect was also reported for *in situ* high-resolution cryo-EM structures of respiratory SCs (50), thus providing additional support for our findings.

We observe that the SC assembly influences the lipid and water dynamics at the protein-lipid interface and perturbs the local dielectric at the membrane plane (Fig. S1E). The dielectric constant (ε⊥) shows a local increase up to 1.5 nm from the membrane before it decays to the aqueous bulk dielectric (Fig. S1C,E). We note that this effect is strongly affected by the lipid type, particularly by the CDL that accumulates around the SC (Fig. 2H, Fig. S2B) and creates a micro-environment, which could enhance a local proton gradient.

Taken together, our combined findings suggest that the SC formation is affected by thermodynamic effects that reduce the molecular strain in the lipid membrane, whilst the perturbed micro-environment also affects the lipid and Q dynamics, as well as the dynamics of the OXPHOS proteins (see below).

### The SC assembly alters the quinone dynamics but does not support substrate channeling

In addition to the phospholipids, the IMM contains 3-5% ubiquinone (Q), which functions as the electron carrier between CI and CIII_2_ in the form of ubiquinol (QH_2_). Our simulations, performed with *ca*. 5% Q/QH_2_ (1:1), suggest that the long isoprenoid tail of ubiquinone (Q_10_) has similar physico-chemical properties as the lipid tails, whilst the Q/QH_2_ headgroup is polar and localizes at the membrane surface, with a flip-flop rate between the leaflets of *ca*. 100-150 ns (Fig. S3A-C). The fast flip-flop motion could support the quinol diffusion from CI to CIII_2_, in which the quinol exits the negatively charged (N-side / matrix side) membrane surface of CI and enters the Q_o_ site of CIII_2_ near the positively charged (P-side / intermembrane space) side of the membrane (see *Discussion*). Interestingly, the Q pool does not accumulate uniformly around the SC. Instead, we observe a *ca*. 3% increase in the local Q concentration near the substrate binding sites of both CI and CIII_2_ relative to the bulk membrane, whilst the immediate OXPHOS surroundings show an overall Q/QH_2_ depletion (Fig. 2I, Fig S2C, Fig S7D,E). This local Q pool could arise from specific interactions between Q and the OXPHOS proteins, *e*.*g*., in subunits ND1 and NDUFA9 of CI that contain several non-polar residues and surface charges that interact with the quinones (Fig. 2M, Fig. S14). In contrast, on the proximal side of CIII_2_, we observe a subtle decrease in the local Q concentration relative to CIII_2_ alone (Fig 2I, Fig S2C, Fig S7D,E), an effect that could arise from a shift in the membrane plane near the Q_o_ site and affect the substrate access to the proximal Q_o_ site (Fig S4, S8). These observations are consistent with the occupied distal Q_o_ site and empty proximal Q_o_ site observed in the cryo-EM structure of the SCI/III_2_ (40).

Overall, while our simulations indicate that quinones accumulate around the OXPHOS proteins, our data do not support substrate channeling (10) (see *Discussion*), as we observe neither a contiguous region of an elevated Q pool between the complexes, nor a directed diffusion pathway of the quinol from CI to CIII_2_ (Fig. 2I, Fig. S3D-I). Moreover, the locally hindered access to the proximal Q_o_ site due to the membrane shift could further hamper the substrate binding into the active site (Fig. S4D-E).

### The SC assembly alters the conformational dynamics of CI and CIII_2_

We next probed how the SC assembly affects the dynamics of the individual OXPHOS proteins based on our atomistic MD simulations of CI, CIII_2_, and the SCI/III_2_. Despite the large (1.5 MDa) SC, we note the SC assembly is stabilized by only a few specific hydrogen-bonding and charged interactions at the interface of NDUFB9/NDUFB4 and UQCRC1, particularly the Arg29^FB4^-Asp260^C1^/Glu259^C1^ ion-paired network that interacts by *ca*. -10 kcal mol^-1^ during the MD simulations (Fig. 3A,C). Indeed, deletion of this Glu258-Asp260 region in UQCRC1 led to a drastic reduction in the stability of SC in mice (29) (see Fig. S11), thus supporting the importance of these interactions in the SC.

**Figure 3.**
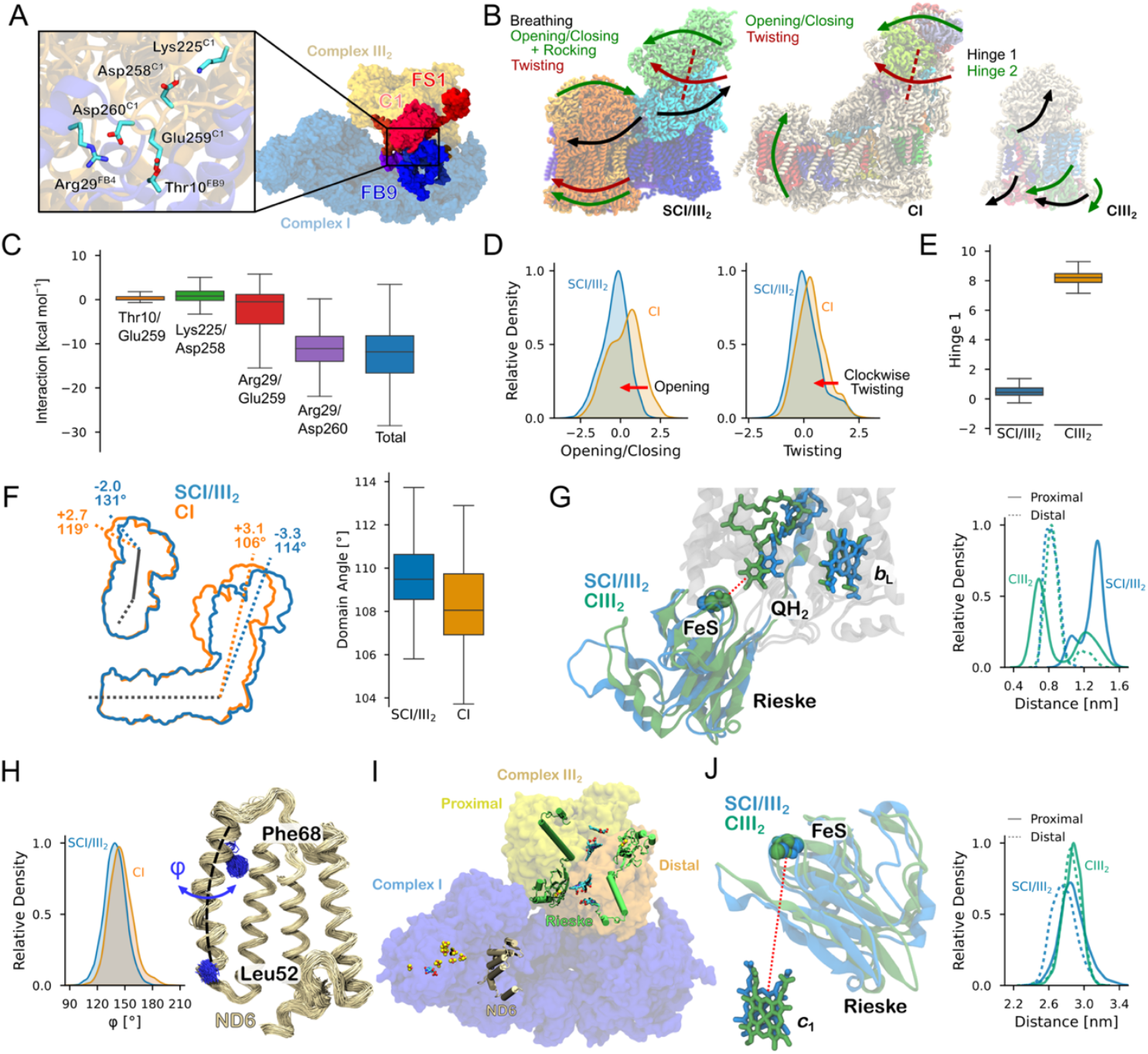
Conformational dynamics of the OXHPOS proteins and the SC. (**A**) Subunits comprising the SC interface. *Inset:* Interaction near the DED-loop. (**B**) Normal modes derived from essential dynamics analysis for the SC (*left*), CI (*middle*), and CIII_2_ (*right*). See SI Appendix, Movies 1-5 for normal modes of the SC, CI, and CIII_2_. (**C**) Decomposition of interaction energies within the SC. (**D**) Distribution of the *opening*/*closing* mode (*left*) and the *twisting* mode (*right*) for the isolated CI (in orange) and the CI within the SC (in *blue*). (**E**) Differences in the dynamics of CIII_2_ for minimum and maximum values of mode 1. (**F**) Distribution of the CI domain angle in the SC (in *blue*) and CI (in *orange*). (**G**,**J**) Conformational changes in the Rieske subunit between SC (in *blue*) and CIII_2_ (in *green*) affect (**G**) the QH_2_ binding in the proximal Q_o_ site and (**J**) the Rieske FeS-heme *c*_1_ distance in the distal monomer. (**H**) Distribution of the dihedral angle in TM3 of ND6. (**I**) Overview of the SC showing the locations of ND6 and Rieske subunits, as well as the proximal and distal protomers of CIII_2_.

Based on essential dynamics analysis (EDA, see *Methods*, Movies S1-S5) that projects out dominant protein motion, we find that the CI and CIII_2_ modules of the SC undergo a back-and-forth *rocking* motion around the interface region, leading to a *breathing* motion between the hydrophilic domains of the OXPHOS complexes on the N-side of the membrane (matrix side) (Fig. 3B, Mode 1, Movie S1), while the second dominant motion (*Mode 2*, Movie S1) couples the *opening*/*closing* motion of CI (cf. also ref (51)) with a *rocking* motion of CIII_2_. The third mode arises from a combination of the *twisting* motion of the hydrophilic domain of CI and a minor *rocking* motion of CIII_2_ (Mode 3, Movie S1), which could influence the membrane accessibility of the Q sites. In this regard, our graph theoretical analysis (Fig. S11C,D) further indicates that ligand binding to Complex I induces a dynamic crosstalk between NDUFA5 and NDUFA10, consistent with previous work (52, 53), and affecting also the motion of UQCRC2 with respect to its surroundings. Taken together, these effects suggest that the dynamics of CI and CIII_2_ show some correlation that could result in allosteric effects, as also indicated based on cryo-EM analysis (40).

We note that the SC assembly induces a subtle conformational change in CI that increases the angle between the hydrophilic and membrane domains (Fig. 3F). This motion is linked to the *active* / *deactive*-transition (A/D) (54), which regulates CI activity, particularly under hypoxic / anoxic conditions, and hinders reverse electron transfer (RET) (54-59) (but *cf*. also ref. (60)). These changes in CI lead to a subtle shift in the conformation of key transmembrane helices (TM) and surrounding conserved loops. In the SCI/III_2_, the TM3 of subunit ND6 undergoes a twist towards the *deactive* state (Fig. 3H, Fig. S13A-D), while the β1-β2 loop of subunit NDUFS2 exhibits lower flexibility in both the *apo* and the QH_2_ bound state relative to CI alone (Fig. S13E-H). These regions are central in modulating the coupling between the proton transport and the electron transfer activities ((61), cf. also Ref. (62)), and the dynamics of these regions could thus have functional consequences. CI also shows a dominant vibrational motion that *twists* the hydrophilic domain relative to the membrane domain (Fig. 3B,D), and samples a wider angle between hydrophilic and membrane domains in the SC, that result in a larger *twisting* angle relative to the isolated CI (Fig. 3F). These alternations suggest that the *opening*/*closing* and *twisting* motions could be sterically hindered by the adjacent CIII_2_ (see Movies S1-S5), and that the *active*/*deactive* (A/D) transition is coupled to the global motions of the SC. Consistent with our previous findings (51), CI shows also another dominant *bending* motion, where the membrane domain rotates relative to the hydrophilic domain within the membrane plane.

CIII_2_ also undergoes dynamical changes upon the SC formation, particularly by enhancing the normal modes connected to the *opening*/*closing* transitions around the iron-sulfur (FeS) Rieske center (Fig. 3B,E) that could affect the electron transfer activity of CIII_2_ (63). In the SCI/III_2_, QH_2_ binds further away from the FeS center in the proximal Q_o_ site, while in the distal site, the FeS center moves closer to the heme *c*_1_ (Fig 3G,I,J). This asymmetry suggests that the global motion of the SC could regulate the ‘preferred’ Q_o_ site for the electron bifurcation process, and possibly favor the electron transfer onwards to the Complex IV (CIV) that resides on the distal side of the CIII_2_ module in the respirasome (SCI/III_2_/IV) (30-32) (cf. also graph theoretical analysis, Fig. S11C,D). Taken together, these effects suggest that the dynamics of CI and CIII_2_ show some correlation that could result in allosteric effects, as also suggested by the recent *in situ* cryo-EM study of mitochondrial SCs (40). In this regard, we find that the ligand state of CI (*apo* or QH_2_) affects the conformational dynamics and the interaction interface of the SCI/CIII_2_ (Fig. S12A-E). This surprising long-range effect is likely to result from the increased flexibility of the *apo* state (Fig. S12G) that, in turn, modulates the interaction at the interface of the SC. Interestingly, similar ligand dependent conformational changes affecting both the CI and CIII_2_ domains of the SC are also supported by recent *in situ* cryo-EM structures of mitochondrial SCs (50).

### Enthalpy-entropy compensation drives SC formation

Our molecular simulations suggest that while the membrane strain provides a thermodynamic driving force for the SC formation, the molecular interactions at the interface of the assembly are essential for enthalpically stabilizing the SC over non-specific protein assemblies (Fig. S12). To assess how the SC formation is affected by the strain effects, protein-protein interactions, and the protein/lipid ratio, we developed a simple statistical-mechanical lattice model of the CI and CIII_2_ diffusion in the IMM (Fig. 4A). To this end, the protein-protein interactions were described by tunable interaction energies (*E*_specific_ and *E*_non-specific_), modeling the specific hydrogen-bond / salt-bridge interactions and the non-specific interaction, while the membrane-protein interactions were tuned by the strain energy (*E*_strain_). Our model suggests that the ratio between the strain energy and specific interactions indeed provides the key driving force for the SC formation, with a small (30%) decrease in *E*_strain_ leading to a *ca*. 60% decrease in the SC population (Fig. 4B). Our model further predicts that the SC population drastically increases at a specific interaction energy threshold (*E*_specific_ < -2 *k*_*B*_*T*) relative to the strain contribution (Fig. 4B), whilst the number of non-specific assemblies (adjacent proteins in non-SC orientations) relative to SCs is determined by the ratio of *E*_specific_ to *E*_non-specific_. With a high membrane strain, we find that an increase in temperature also favors the formation of SC assemblies, as it is entropically favored to reduce the overall number of ordered (“strained”) lipid molecules around the OXPHOS proteins (Fig. S15). Similarly, a high *protein*-to-*lipid* ratio significantly increases the population of SCs, suggesting that a high protein packing in IMMs favors the SC formation, whilst the strength of specific interactions determines the relative population of non-specific assemblies in crowded environments. Taken together, our lattice model, despite its simplicity, captures and validates key features observed in our molecular simulations and supports that the SC formation is affected by both enthalpic and entropic effects.

**Figure 4.**
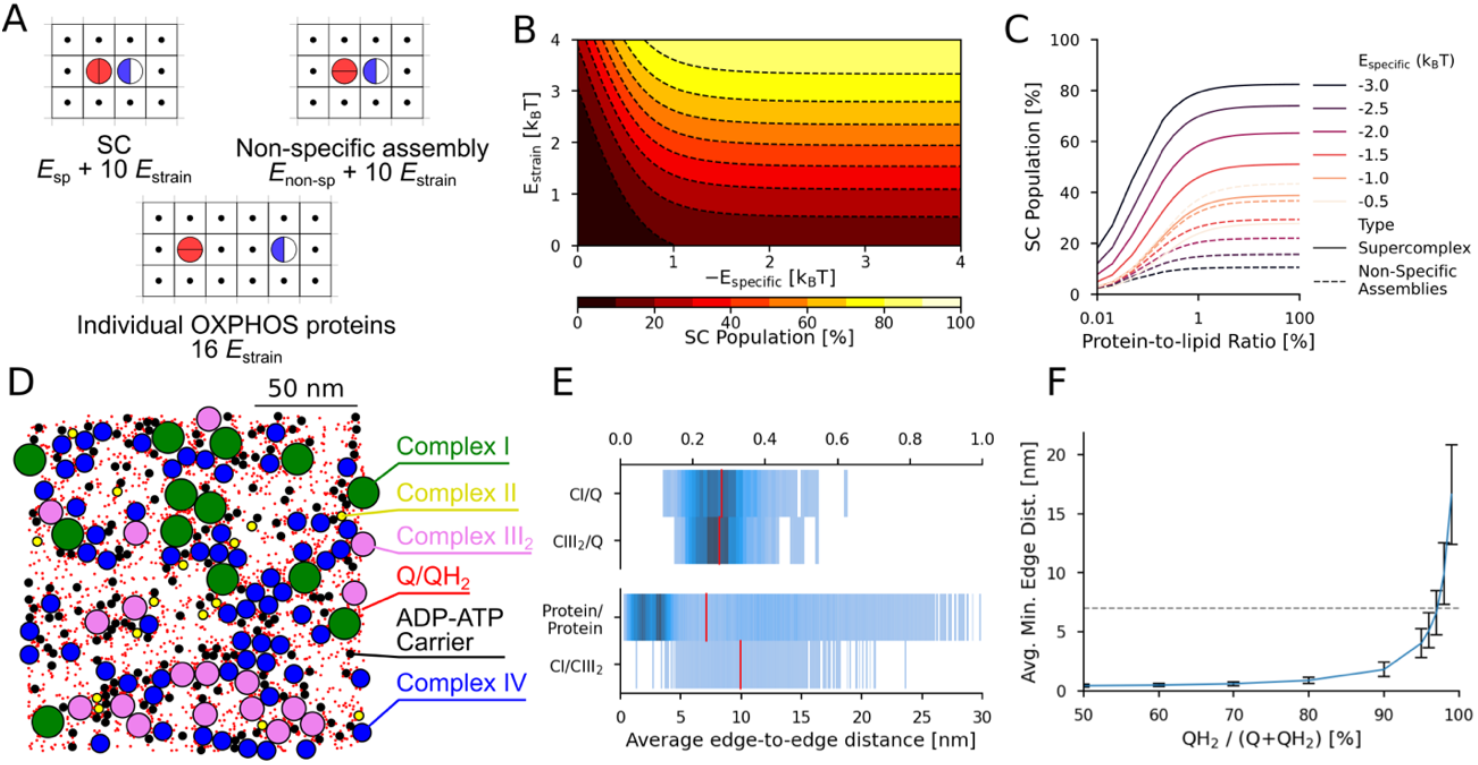
Lattice model of SC formation and crowding effects in the IMM. (**A**) Possible specific and non-specific interactions between CI (blue/white) and CIII_2_ (red) and their respective energies in the lattice model. (**B**) SC population as a function of the specific interaction energy (*E*_specific_) and molecular strain (*E*_strain_). (**C**) Specific and non-specific assemblies as a function of the protein-lipid ratio with varying specific interaction energies (*E*_specific_). (**D**) Representative protein arrangement in crowded IMMs. Average *edge-to-edge* distance distributions and nearest neighbor distance (red line) for specific protein-Q/protein contacts. Nearest neighbor distance between Q and CI as a function of the Q/QH_2_ ratio. The dashed line indicates the 7 nm distance between active sites in the SC.

## Discussion

We have shown here that the respiratory chain complexes perturb the IMM by affecting the local membrane dynamics. The perturbed thickness and alteration in the lipid dynamics lead to an energetic penalty, which can be related to molecular strain effects, as suggested by the changes of both the internal energy of lipid and their interaction with the surroundings (Fig. S2, S5, S6), which are likely to be of enthalpic origin. However, lipid binding to the OXPHOS complex also results in a reduction in the translational and rotational motion of the lipids and quinone (Fig. S9, S10), which could result in entropic changes. The strain effects are therefore likely to arise from a combination of enthalpic and entropic effects. In this regard, we suggest that the SC assemblies form by condensation of local high-entropic membrane regions around the OXPHOS proteins (Fig. 5A). However, as the entropic effects also favor the formation of non-specific protein assemblies, unique interactions, such as the ion-paired network around UQCRC1 and NDUFB9/NDUFB4 (Fig. 3A), are likely to enthalpically stabilize the SC assemblies. The suggested principles show similarities to the hydrophobic effect driving protein folding by condensation of locally ordered water clusters around unfolded protein patches (64).

**Figure 5.**
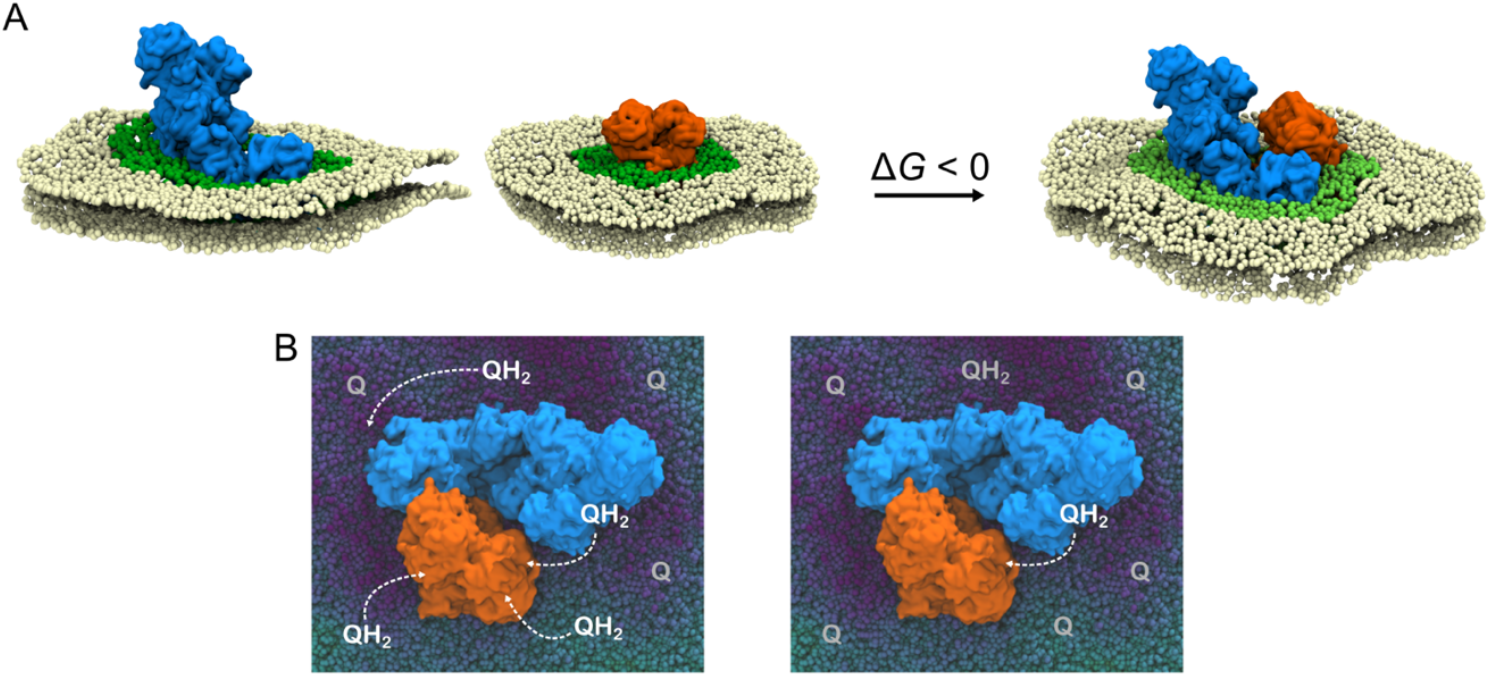
Proposed thermodynamic and functional effects of SCs. **(A)** The individual OXPHOS complexes, CI and CIII_2_ induce prominent molecular strain in the surrounding lipid membrane (dark green area) due to a hydrophobic mismatch between the protein and the membrane. Condensation of locally strained membrane patches entropically drives the SC formation, and leads to an overall reduction of the membrane strain (light green area around the SC), favoring the accumulation of cardiolipin around the SC. **(B)** During normal respiratory conditions, each OXPHOS complex is surrounded by multiple (ca. 6) quinol/quinone molecules that can act as substrates for the proteins. At limiting QH_2_ concentration, the quinol diffusion between CI and CIII_2_ becomes rate-limiting, and leads to a kinetic advantage of the SC (see also Fig. 4F).

We note that the magnitude of the estimated bending energies (∼10^2^ kcal mol^−1^) (Table 1), while seemingly high at first glance, falls within the range expected for large-scale membrane deformation processes induced by large multi-domain proteins. For example, the Piezo mechanosensitive channel performs roughly 150*k*_B_*T* (≈ 90 kcal mol^−1^) of work to bend the bilayer into its dome-like shape (65). Comparable energies have also been estimated for the nucleation of small membrane pores (66), while vesicle formation typically requires bending energies on the order of 300 kcal mol^−1^, largely independent of vesicle size (67). When normalized by the affected membrane area (∼1000 nm^2^), these values correspond to an energy density of approximately 0.1 kcal mol^−1^ nm^−2^, which places our estimates within a biophysically reasonable regime. Notably, cryo-EM structures of supercomplexes shows that such assemblies can impose significant curvature on the surrounding bilayer (36, 50, 68), supporting the notion that respiratory chain organization is closely coupled to local membrane deformation. Nevertheless, we expect that the absolute deformation energies may be overestimated, as the continuum Helfrich model neglects molecular-level effects such as lipid tilt and local rearrangements, which can partially relax curvature stresses and reduce the effective bending penalty near protein–membrane interfaces (69, 70).

We further probed the thermodynamic effects underlying the SC formation by developing a 2D lattice model. Despite its simplicity, the model supports that the SC stability is determined by a delicate balance between membrane strain effects and specific protein-protein interactions, but also strongly affected by the protein concentration and temperature effects (6Fig. 4B). Moreno-Loshuertos *et al*. (71) recently suggested that elevated temperatures (>43°C) may indeed disrupt SCs, although the stability of the individual OXPHOS proteins was also decreased in the studied conditions. As the enthalpic effects are of electrostatic origin, the SC stability could also be sensitive to the ionic strength and the PMF, which in turn depends on the metabolic state of the mitochondria.

At the molecular level, the SC formation leads to an accumulation of CDL at the protein-membrane interface (Fig. 2N), as well as a local Q / QH_2_ pool near the substrate channel of CI and CIII_2_ (Fig. 2M). We find that CDL prefers thinner membranes relative to the neutral phospholipids (PE/PC, Fig. S5E), and could thus partially compensate for the hydrophobic mismatch between the OXPHOS proteins and the membrane (Fig. S1). The Q diffusion is affected by both specific interactions with OXPHOS proteins, as well as the local membrane thickness (Fig. 2I). The CDL around the SC is thus likely to have both structural and functional consequences, consistent with its effect on the activity and dynamics of several membrane proteins ((72-75), *cf*. also Ref. (51)). In this regard, CDL was suggested to enhance the substrate dynamics within the Q-tunnel of CI that requires a *twisting*-*bending* motion around the membrane and hydrophilic domains (51), whilst destabilization of CDL-binding sites has indeed been shown to disrupt SCs, *e*.*g*., the SCIII_2_IV_1_ in yeast (12). Moreover, defects in the CDL synthesis, *e*.*g*., in the Barth syndrome, (76), result in the disassembly of SC, indirectly supporting the involvement of CDL as a “SC glue”. In this regard, electrostatic effects arising from the negatively charged cardiolipin headgroup could play an important role in the interaction of the OXPHOS complexes. While CDL was modeled here in the singly anionic charged state (but cf. Fig. S5E), we note that the local electrostatic environment could tune their p*K*_a_ that result in protonation changes of the lipid, consistent with its function as a proton collecting antenna (77).

In addition to the changes in the membrane properties, we observed that the SC formation modulates the conformational dynamics of the individual OXPHOS proteins, especially the large-scale *bending*-*twisting* motion of CI, the conformation of individual TM helices and conserved loops around proton channels in CI, as well as the motion of the Rieske domain in CIII_2_ (Fig. 3, Fig. S12). The dynamics of these regions are likely to module the activity of the OXPHOS proteins, *e*.*g*., the A/D transition of CI (59, 78) that regulate the ΔpH-driven quinol oxidation and reverse electron transfer (79, 80). Indeed, blocking loop motions surrounding these regions inhibits the proton pumping activity of CI (81), whilst the perturbed motion of the Rieske domain could modulate the electron bifurcation in CIII_2_ and subsequent electron transfer to CIV. In this regard, we suggest the preferred QH_2_ binding in the distal Q_o_ site has functional implications for the respirasome (SCI/III_2_/IV), where the CIV module is located on the distal side of the CIII_2_ protomer. The changes in the conformational dynamics upon SC formation may thus affect ROS production (17, 82) via the A/D transition of CI (59), although it should be noted that no differences in ROS generation were observed for mice unable to form SCs relative to WT mice (with *ca*. 75% SCs) under normal conditions (29). It is possible that differences occur only under more strained conditions, *e*.*g*., in hypoxia or together with disease-related mutations in the OXPHOS proteins, where the SCs could become functionally more important (see below).

To understand how SCs could influence the charge currents in the IMMs, we note that the local increase of the quinol concentration near the active site of CI, and the local decreased quinol pool around the proximal Q_o_ site of CIII_2_ could create a substrate gradient (∇*c*) and affect the quinol flux between CI and CIII_2_, *J*_CI→CIII_ = -*D*_*mem*_∇*c* (*D*_*mem*_ – Q/QH_2_ diffusion constant) – if the quinol concentration is rate-limiting for the function of CI or CIII_2_ (but see below). The 2D-diffusion time for the quinol, *τ* = <*r*^2^>/4*D*_*mem*_, between the OXPHOS complexes depends on the effective protein-protein distance (*r*), which can be estimated from the protein packing density (cf. (7, 34, 83-87)). Using the experimental protein copy numbers in IMM (cf. Refs. (7, 34, 83-87) and Extended SI Text, Table S5), we obtain average *edge-to-edge* distance between CI and CIII_2_ of around 12 nm (Fig. 4E, *cf*. also (7)), which can be compared to the *edge-to-edge* distance of *ca*. 7 nm / 12 nm between the Q tunnel of CI and the proximal / distal Q_o_ sites of CIII_2_ within the SC. The reduced diffusion distance could thus provide a subtle rate enhancement (*τ*_SC_(*r*=7 nm)/*τ*_nonSC_(*r*=12 nm)∼0.3) in favor of the SCs under substrate-limited conditions (Fig. 4E-F). We expect that the small variations in the lateral quinol diffusion (0.4 nm ns^-1^ near CI, 0.8 nm ns^-1^ in bulk) around the SC (Fig S3D-I, Fig S9) could favor the diffusion along the membrane patches, although this diffusion is also affected by non-specific collisions. However, due to the overall 3-5% Q / QH_2_ concentration in IMMs, we estimate that each OXPHOS protein is surrounded by around 6 quinone / quinol species depending on the respiratory conditions (Fig. 5, Extended SI Text), and leading to a small (0.3 nm) nearest neighbor distance between Q and CI/CIII_2_ (Fig. 4E, see SI Appendix). This implies that each OXPHOS protein has a saturated substrate Q pool within its “reaction sphere” upon high charge flux conditions (here 50% reduced Q pool). In other words, the CIII_2_ node of the SC does not need to rely on the quinol generated at the CI node, but can instead oxidize a quinol molecule from the surrounding Q pool. In this regard, Hirst and co-workers (88) found that the alternative oxidase (AOX) co-reconstituted into liposomes outcompetes the quinol re-oxidation rate of the SCI/III_2_, suggesting that the diffusion of the Q-species between the OXPHOS complexes is not rate-limiting under the studied conditions. However, at low QH_2_/Q ratios (<5% in the model), the minimal nearest neighbor distance between the Q pool and CI/CIII_2_ drastically increases (Fig. 4D-F) that in turn leads to a kinetic preference for the SC (Fig. 4D).

At the microscopic level, the SC influences the lipid and water dynamics at the protein-lipid interface and the physico-chemical properties of the IMM, including its local dielectric properties of the membrane. It is possible that such local micro-environments have functional implications that could affect the proton conduction along the membrane, with important bioenergetic consequences. Taken together, our combined findings suggest that SC forms as a result of a complex interplay between molecular interactions and membrane strain effects that control the functional dynamics of the OXPHOS proteins.

## Materials and Methods

### Atomistic MD Simulations

Atomistic molecular dynamics (aMD) simulations of the ovine Complex I (CI), Complex III_2_ (CIII_2_), and the I-III_2_ supercomplex (SCI/III_2_) were performed in a POPC:POPE:cardiolipin (2:2:1) membrane containing 5 mol% QH_2_ / Q (1:1 ratio). Cardiolipin was modeled as tetraoleoyl cardiolipin (18:1,18:1,18:1,18:1) with a headgroup modeled in a singly protonated state (with Q_tot_=-1). The system was solvated with TIP3P water molecules and 150 mM NaCl. The simulations were performed in different ligand states (*apo*, Q/QH_2_ bound, Table S1) at *T*=310 K and p=1 bar using NAMD2.14 (89) with a 2 fs integration timestep and long-range electrostatic interactions treated using the Particle Mesh Ewald approach. During construction of the simulation setups, it was carefully considered that no leaflet introduced higher lipid densities that could result in artificial displacement effects. The systems comprised 0.8-1.65 million atoms and were modeled using the CHARMM36 force field (90) in combination with in-house DFT-based parameters (91) of the co-factors. Protonation states were established based on electrostatic calculations with Monte Carlo sampling techniques using APBS/Karlsberg+ (92-94).

Molecular dynamics (MD) simulations of the ovine CI were conducted based on a cryo-EM structure (PDB ID: 6ZKC (54)), with simulations performed for both the quinol-bound and *apo* forms. Unresolved regions in the supernumerary subunits NDUFA7 and NDUFB6 were modeled using ColabFold (95). The system was equilibrated for 100 ns in the QH_2_-bound state, which was then used to propagate both states for 2×0.5 µs each.

The ovine Complex III_2_ was modeled based on the cryo-EM structure (PDB ID: 6Q9E (40)), with simulations also performed in different ligand states (Table S1). After equilibration of the system for 100 ns with the distal and proximal Q sites occupied, the system was propagated in each state for 2×0.5 µs.

A fully atomistic model of the SC was constructed based on the high-resolution structures of CI and CIII_2_, that were merged into the experimental structure of the supercomplex (PDB ID: 6QBX (40)). After equilibration of the system for 100 ns with all Q sites occupied in CI and CIII_2_, the complete system was propagated for 2×0.5 µs in each state (see Table S1).

Membrane systems with 1:1 POPC/POPE or CDL were constructed using CharmmGUI with a membrane area of 80×80 Å^2^, a hydration layer of 45 Å, and NaCl or KCl concentrations of 150 mM. See Table S2 for a detailed description of the membrane compositions. See the SI Appendix for a detailed description of the simulations and analysis.

### Coarse-grained MD simulations

Coarse-grained MD (cgMD) simulation models of the ovine CI, CIII_2_, and the SCI_1_/III_2_ were created based on the atomistic models using the MARTINI3 force field (96) and Gromacs (97). All simulations were constructed with identical amounts of lipid molecules, a 2:2:1 POPC:POPE:cardiolipin ratio, and a 3 mol% mixture of quinone/quinol embedded in a simulation system with dimensions 47×47×31 nm^3^ (for the SC, see Table S1). All simulations were carried out at 310 K in an *NPT* ensemble using the velocity rescaling thermostat (98, 99), the Parrinello-Rahman barostat (100) and a 20 fs timestep using GROMACS (97). The protein structures were stabilized with elastic networks on backbone beads using a cutoff distance of 0.9 nm with a force constant of 500 kJ mol^−1^ nm^−2^. The elastic network was also applied between residues of different subunits, but not between CI and CIII_2_, whilst cgMD parameters for all cofactors were also developed. Two replicas were run for CI and CIII_2_ (2×50 µs each), and for the SCI/III_2_ (2×50 µs), as well as 23 µs cgMD simulations of the membrane with the same number of lipid and Q/QH_2_ molecules as in the protein simulations. Additional 5 µs cgMD simulations of the membrane systems, as well as for all protein models, were performed with longer lipids (0.44 nm, 0.50 nm, and 0.53 nm instead of 0.47 nm).

### Lattice model of SC formation

A lattice model of the CI and CIII_2_ was constructed (Fig. 4A,B) by modeling the OXPHOS proteins in unique grid positions on a 2D *N*×*N* lattice, with *N*=[6,15]. Depending on the relative orientation, the protein-protein interaction was described by specific interactions (giving rise to the energetic contribution *E*_specific_ < 0) and non-specific interactions (*E*_non-specific_ > 0), whereas the membrane-protein interaction determined the strain energy of the membrane (*E*_strain_), based on the number of neighboring “lipid” occupied grids that are in contact with proteins (Fig. 4A). The interaction between the lipids was indirectly accounted for by the background energy of the model. The proteins could occupy four unique orientations on a grid (*g*_*i*_=[*North, East, South, West*]). The states and their respective energies that the system can visit are summarized in Table S6.

The total Hamiltonian of the lattice model can be written as,

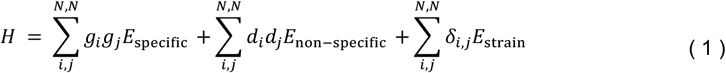

With

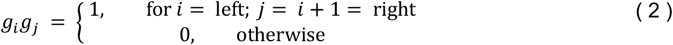

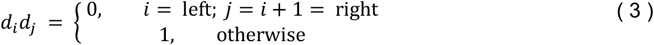

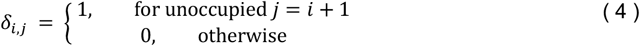

The *g*_i_*g*_j_ -term assigns a specific energy contribution when the OXPHOS complexes are in adjacent lattice points only in a correct orientation (modeling a specific non-covalent interaction between the complexes such as the Arg29^FB4^-Asp260^C1^/Glu259^C1^ interaction between CI and CIII_2_). The *d*_i_*d*_j_ -term assigns a non-specific interaction for the OXPHOS complexes when they are in adjacent lattice points, but in a “wrong” orientation relative to each other to form a specific interaction. The δ_ij_ term introduces a strain into all lattice points surrounding an OXPHOS complex, mimicking the local membrane perturbation effects observed in our molecular simulations.

This leads to the partition function,

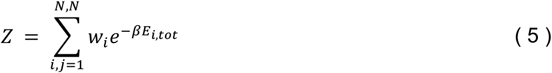

where *w*_i_ is the degeneracy of the state, modeling that the SC and OXPHOS proteins can reside at any lattice position of the membrane, and where *β*=1/*k*_B_*T* (*k*_B_, Boltzmann’s constant; *T*, temperature). The probability of a given state *i* was calculated as,

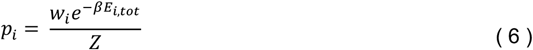

with the free energy (*G*) defined as,

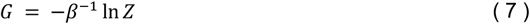

The conformational landscape was sampled by Monte Carlo (MC) using 10^7^ MC iterations with 100 replicas (see SI Appendix). Temperature effects were modeled by varying *β*, and the effect of different *protein*-to-*lipid* ratios by increasing the grid area. The simulation details can be found in Table S7.

### Statistical model of the membrane distribution in the IMM

Based on the experimental protein copy numbers (*cf*. Ref (7)) and average protein areas, the proteins in the IMM were modeled as randomly positioned circles on a membrane square with a side length of 163 nm (see Table S5). The system was minimized according to the Hamiltonian,

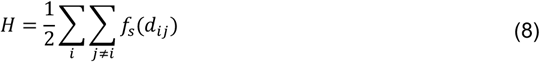

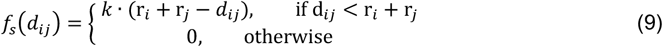

where *d*_ij_ is the distance between two proteins *i* and *j* with respective radii, *r*_*i*_ and *r*_*j*_, while the force constant *k*, were introduced to avoid steric clashes and set to 1 kcal mol^-1^ nm^-1^. This Hamiltonian corresponds to a two-dimensional system consisting of rigid, non-deformable circular particles.

## Acknowledgments

This work was supported by grants from the European Research Council (ERC) under the European Union’s Horizon 2020 research and innovation program/grant agreement 715311, the Swedish Research Council (VR), and the Knut and Alice Wallenberg foundation. (2019.0251, 2019.0043, 2024.0220), and the Göran Gustafsson Foundation for Research in Natural Sciences and Medicine. We are thankful for computing time provided by the Partnership for Advanced Computing in Europe (PRACE project: pr127) to access Piz Daint hosted by the Swiss National Supercomputing Center (CSCS). This work was also supported by the National Academic Infrastructure for Supercomputing in Sweden (NAISS 2025/1-33, 2025/6-165, 2024/1-28, 2023/1-31, 2023/6-128) and the Swedish National Infrastructure for Computing (SNIC 2022/1-29, 2022/6-190).

## Notes

### Competing Interest Statement

The authors have declared no competing interest.

### Summary of Updates

The magnitude of the membrane curvature term is discussed in the revised manuscript.

