## Supporting Information for "Protein-Induced Membrane Strain Drives Supercomplex Formation"

**This PDF file includes:**

Supporting methods
Figures S1 to S15
Tables S1 to S7
Movies S1 to S5
SI References

**Other supporting materials for this manuscript include:**

Movies S1 to S5

**Supporting Methods**

**Estimation of membrane strain.** The potential of mean force of the membrane thickness profile ($z$) was calculated based on,

$$G_{\mathrm{memb}}(z) = -k_B T \ln \rho(z) \tag{1}$$

where $z$ is the membrane thickness and $\rho$ is the probability distribution of the membrane thickness. The membrane strain induced by the protein was calculated from the cgMD simulations of the different systems as (see *Methods*),

$$G_{\mathrm{strain}}^{\mathrm{protein}} = \int \rho^{\mathrm{protein}}(z) G_{\mathrm{memb}}(z)\, \mathrm{d}z \tag{2}$$

To obtain detailed atomistic insight into the strain effects, the local lipid strain was quantified by energy decomposition analysis, by comparing the internal lipid energies to the average lipid energy from a membrane simulation (Fig. S5). The simulation data were averaged over 690,000 datapoints from 1.85 µs MD simulations of systems A1,7,9 and M4 (Table S1-S2).

**Estimation of dielectric profiles**

The dielectric constant (ε) was determined from the local variance of the dipole moment at a given position $\boldsymbol{r}_p$ using the Kirkwood-Fröhlich equation (1, 2),

$$\varepsilon = 1 + \frac{\mathrm{var}\, \boldsymbol{M}}{3\varepsilon_0 k_B T V} \tag{3}$$

with the local dipole moment was calculated from the sum of the charge-weighted positions of all atoms within 0.2 nm of $\boldsymbol{r}_p$,

$$\boldsymbol{M}(\boldsymbol{r}_p, t) = \sum_{i,\ |r_i - r_p| < 0.2\ nm} q_i \boldsymbol{r}_i \tag{4}$$

where $\varepsilon_0$ is the vacuum permittivity, $k_B$ is the Boltzmann constant by, $T$ is the temperature, and $V$ is the volume of the selection sphere. The calculations were performed by averaging the total **M** over fixed $z$ values from the membrane plane. Note that this treatment differs from extraction of radial and axial contributions of the dielectric tensor, as developed by Netz and co-workers (cf. Ref. (3) and refs therein) that requires a more elaborate treatment, which is outside the scope of the present work.

**Analysis of lipid chain *end-to-end* length**

To probe the protein-induced deformation effect of the membrane, the membrane curvature ($H$), and the *end-to-end* distance between the lipid chains, were computed based on aMD and cgMD simulations. The lipid chain length was computed from simulations A1-A6 and C1 based on the first and last carbon atoms of each lipid chain. For example, the *end-to-end* length of a cardiolipin chain was determined as the distance between atom "CA1" and atom "CA18".

**Membrane Curvature and Deformation Energy**

The local mean curvature of the membrane midplane was computed using the Helfrich model (4,5) by approximating the membrane surface as a height function $Z(x,y)$, defined as the average location of the N-side and P-side leaflets at each grid point. Based on this, the mean curvature $H(x,y)$ was calculated as,

$$H(x,y) = \frac{(1+Z_x^2)Z_{yy} + (1+Z_y^2)Z_{xx} + 2Z_xZ_yZ_{xy}}{2(1+Z_x^2+Z_y^2)^{\frac{3}{2}}} \tag{5}$$

where the derivatives are defined as $Z_{\mathrm{i}} = \frac{\partial Z}{\partial \mathrm{i}}$ and $Z_{ij} = \frac{\partial^2 Z}{\partial i\, \partial j}$ .

The thickness deformation energy was computed from the local thickness $d(x,y)$ relative to a reference thickness distribution $F$(d), derived from membrane-only simulations, and converted to a free energy profile via Boltzmann inversion. At each grid point, the $F(d)$ was summed over the grid,

$$G_{thick} = \sum_{x,y} F\big(d(x,y)\big)\, \Delta A \tag{6}$$

The bending deformation energy was computed from the mean curvature field $H(x,y)$, assuming a constant bilayer bending modulus κ (taken as $20k_bT$ = 11.85 kcal mol$^{-1}$ (6)):

$$G_{curv} = \frac{1}{2}\kappa \sum_{x,y} (2H(x,y))^2\, \Delta A \tag{7}$$

where $\Delta A$ is the area of the grid cell.

The thickness and curvature fields were obtained by projecting the coarse-grained MD trajectories (one frame per ns) onto a 2D-grid with a resolution of 0.5 nm. Grid points with low occupancy were down-weighted to mitigate noise. More specifically, points with counts below 50% of the median grid count were scaled linearly by their relative count value. To focus the analysis on the region around the protein–membrane interface, only grid points within a radius of 20 nm from the center of the complex were included in the energy calculations. Energies were normalized to an effective membrane area of 1000 nm$^2$ to facilitate the comparison between systems. Bootstrapping with resampling over frames was performed to estimate the standard deviations of $G_{thick}$ and $G_{curv}$.

We find that $G_{curve}$ converges slowly due to its sensitivity to local derivatives and the small grid size required to resolve the curvature contribution near the protein. Consequently, tens of microseconds of simulations were necessary to obtain well-converged estimates of the curvature energy.

**Spatial integration of cryo-EM maps.** The properties of the local membrane environment were estimated based on a high-resolution structure of Complex I (PDB ID: 6RFR, EMD-4873) and the associated cryo-EM map (7). To this end, we first subtracted the cryo-EM density close to the resolved protein structure, whilst the density vertical to the membrane plane was approximated as a double sigmoid function. To identify the membrane regions, we used a neural network model (multi-layer perceptron classifier) with 3×100 hidden layers that we trained to differentiate between double sigmoid-type regions and regions with different electron density distributions. To this end, we created 20,000 random double-sigmoid distributions with random errors as well as 120,000 of various random distributions. We used 25% of the data as test data, resulting in an accuracy score of 97%.

**Estimation of effective Q/QH$_2$ concentrations.** The IMM has an area of around 16,905 nm$^2$ and it comprises 48,300 lipid molecules (Table S5). For a 1% Q/QH$_2$ concentration, this implies 483 Q/QH$_2$ molecules. Based on protein copy numbers and the structure the membrane proteins, the IMM comprises 291 proteins with a total area of 14,586 nm$^2$ (Table S5), thus suggesting that two mitochondrial proteins occupy an average area of 216.4 nm$^2$. For a square membrane patch, this would lead to an edge of $R$ = 14.7 nm, and an average protein-protein distance of,

$$
\begin{aligned}
\frac{\langle r_{\mathrm{p-p}} \rangle}{R} &= \int_0^1\int_0^1\int_0^1\int_0^1 \sqrt{(x_1^2 - x_2^2)^2 + (y_1^2 - y_2^2)^2}\,\mathrm{d}x_1\mathrm{d}y_1\mathrm{d}x_2\mathrm{d}y_2 \\
&= 4\int_0^1\int_0^1 \sqrt{x^2 + y^2}\,\mathrm{d}x\mathrm{d}y \\
&= 8\int_0^{\frac{\pi}{4}}\int_0^{\frac{1}{\cos\theta}} (1 - r\ \sin\theta)(1 - r\ \cos\theta) r^2\, dr d\theta \\
&= \frac{1}{15}\left[2 + \sqrt{2} + 5\ln\left(\sqrt{2} + 2\right)\right] \approx 0.52
\end{aligned}
\tag{5}
$$

or $\langle r_{\text{p-p}} \rangle$ = 7.6 nm. For the same membrane patch, we thus have an average of 216.4 nm$^2$ × ((48,300×0.01)/16,906.0 nm$^2$) = 6.2 Q/$QH_2$ molecules.

**Multiple Sequence Alignment (MSA).** Sequences of mammalian species were aligned using the ClustalOmega (8) command line interface of Biopython (9). Sequences of selected species were visualized using Jalview (10).

**Allosteric Network Analysis.** Interactions between amino acid residues were modeled as an interaction graph, where each residue was represented by a vertex. Two nodes were connected by an edge, if the C$\alpha$ atoms of the corresponding amino acid residues were closer than 7.5 Å for more than 50% of the frames of simulations A1-A6 (time step of frames: 1 ns). (11) This analysis was carried out for the aMD simulations of the supercomplex, analyzing differences between the Q bound and *apo* states (simulations A1+A2+A3 vs. A4+A5+A6).

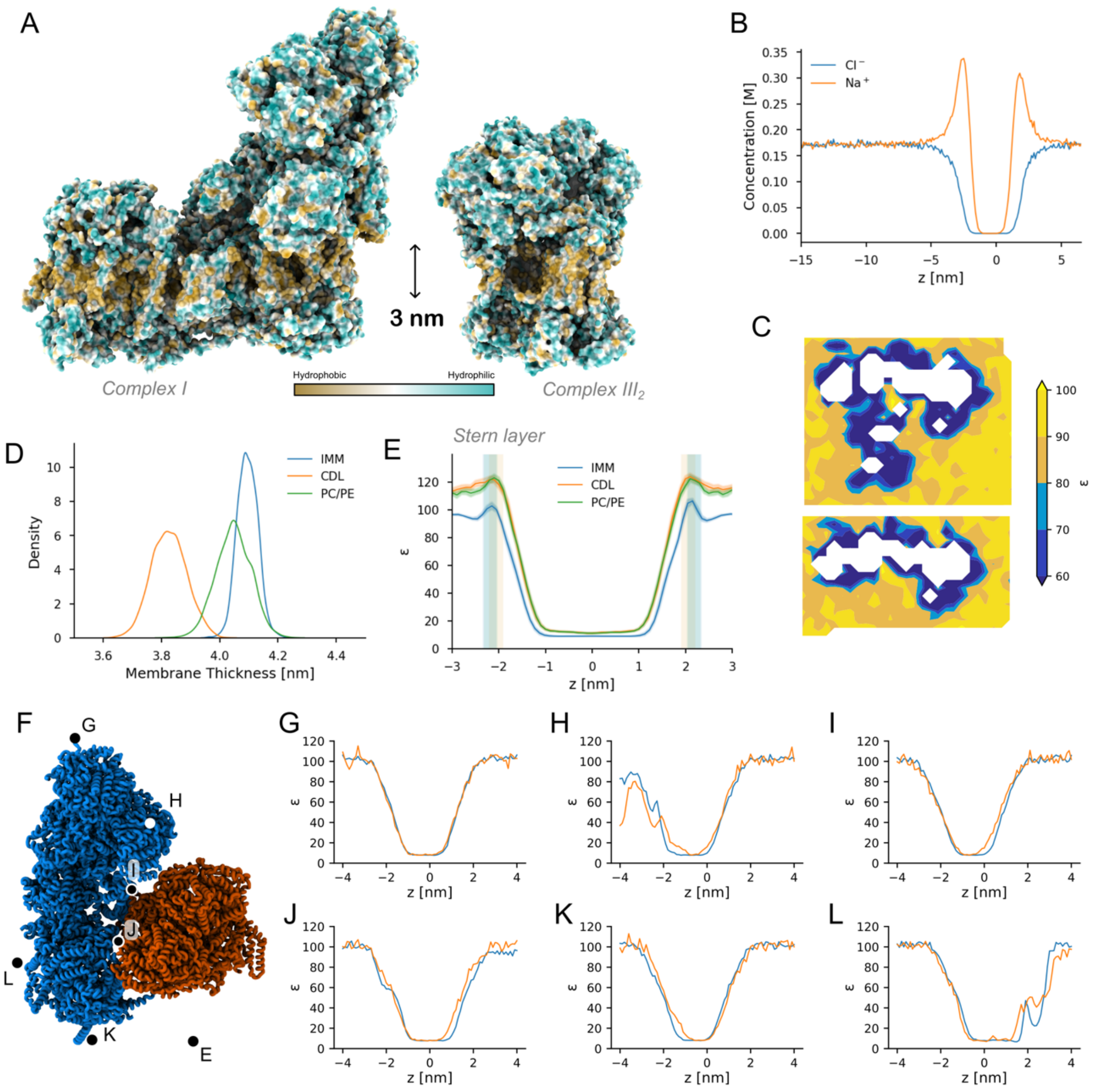


**Fig. S1.** Analysis of membrane properties. Mismatch of the hydrophobic region of CI and CIII$_2$ relative to the lipid membrane. (**A**) The hydrophobic region in CI (*left*) and CIII$_2$ (*right*), with hydrophobic amino acids in gold, and hydrophilic amino acids in cyan. The maps were created using ChimeraX. (**B**) Ion distribution as a function of *z* position in a protein-free slab of simulation S1. (**C**) Dielectric constant (at *z*=0.5 nm from the membrane-water interface) around the SC (*left*) and CI (*right*), with the SC leading to larger membrane surface with a perturbed ε. (**D**) Distribution of the membrane thickness in a POPC/POPE/CDL/$QH_2$ membrane and a CDL membrane. (**E**) Dielectric profile (ε) computed perpendicular to the membrane plane for a POPC/POPE and a pure CDL membrane. The Stern layer, characterized by the local increase in the ε, extends *ca.* 1.5 nm from the membrane plane and is affected by the lipid composition. (**F**) Position of sampled dielectric profiles shown in (**G**) to (**L**).

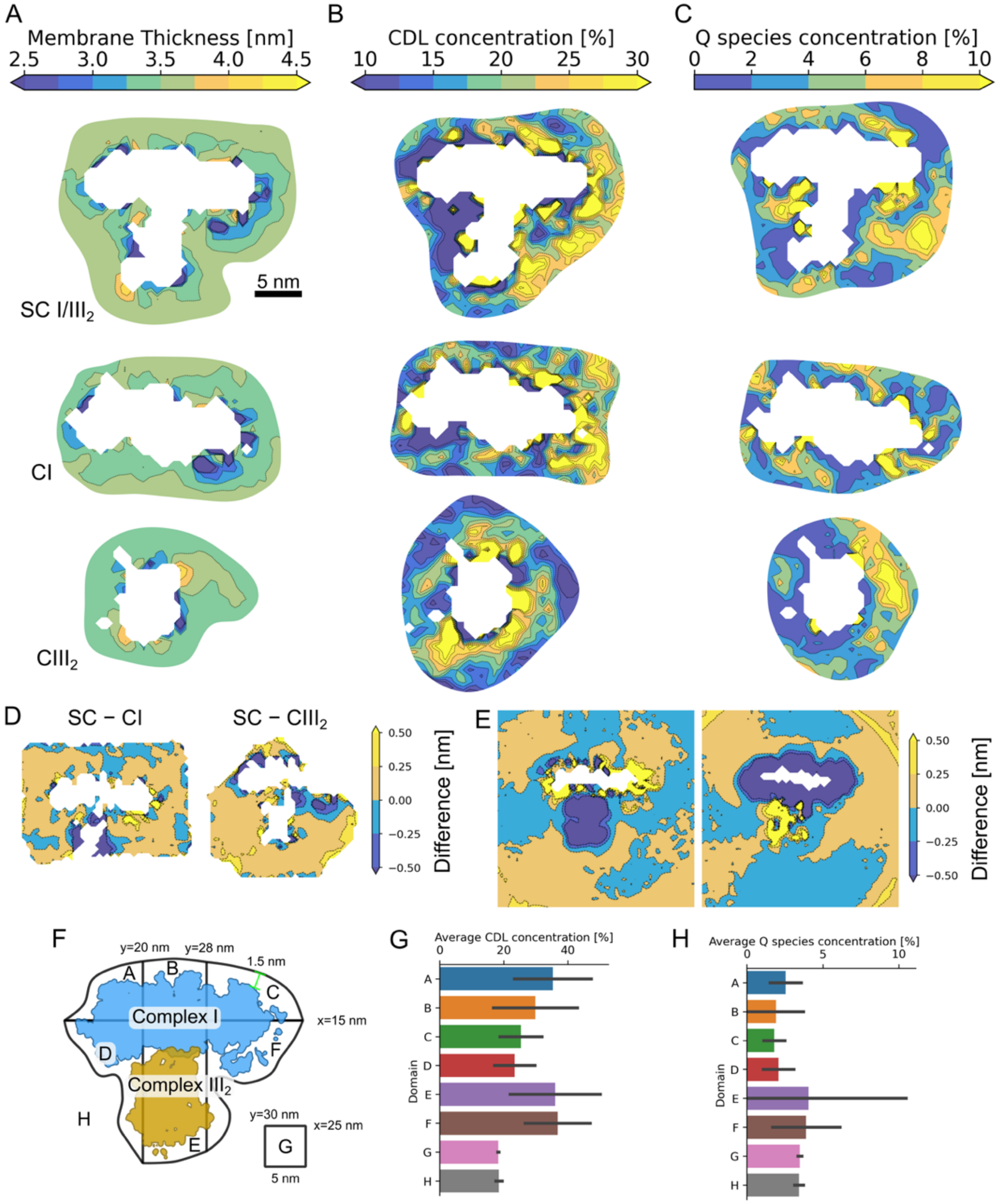


**Fig. S2.** Concentration of lipids and quinones, and analysis of membrane thickness in aMD simulations. (**A**) Local membrane thickness in the SCI/CIII$_2$ (*top*), CI (*middle*), and CIII$_2$ (*bottom*). (**B**) Local cardiolipin concentration. (**C**) Local $QH_2$ concentration. (**D, E**) Difference in the membrane thickness around the SC relative to CI (*left*) or relative to CIII$_2$ (*right*) from (**D**) aMD and (**E**) cgMD. **(F, G, H)** Error bars for the local CDL and Q/$QH_2$ enrichment (data presented in Fig. 2G,H) were estimated by averaging over the membrane in the vicinity of the protein (within 1.5 nm distance) divided into six domains (see panel F). The average concentrations for each domain and the calculated errors of (**G**) CDL and (**H**) Q/$QH_2$.

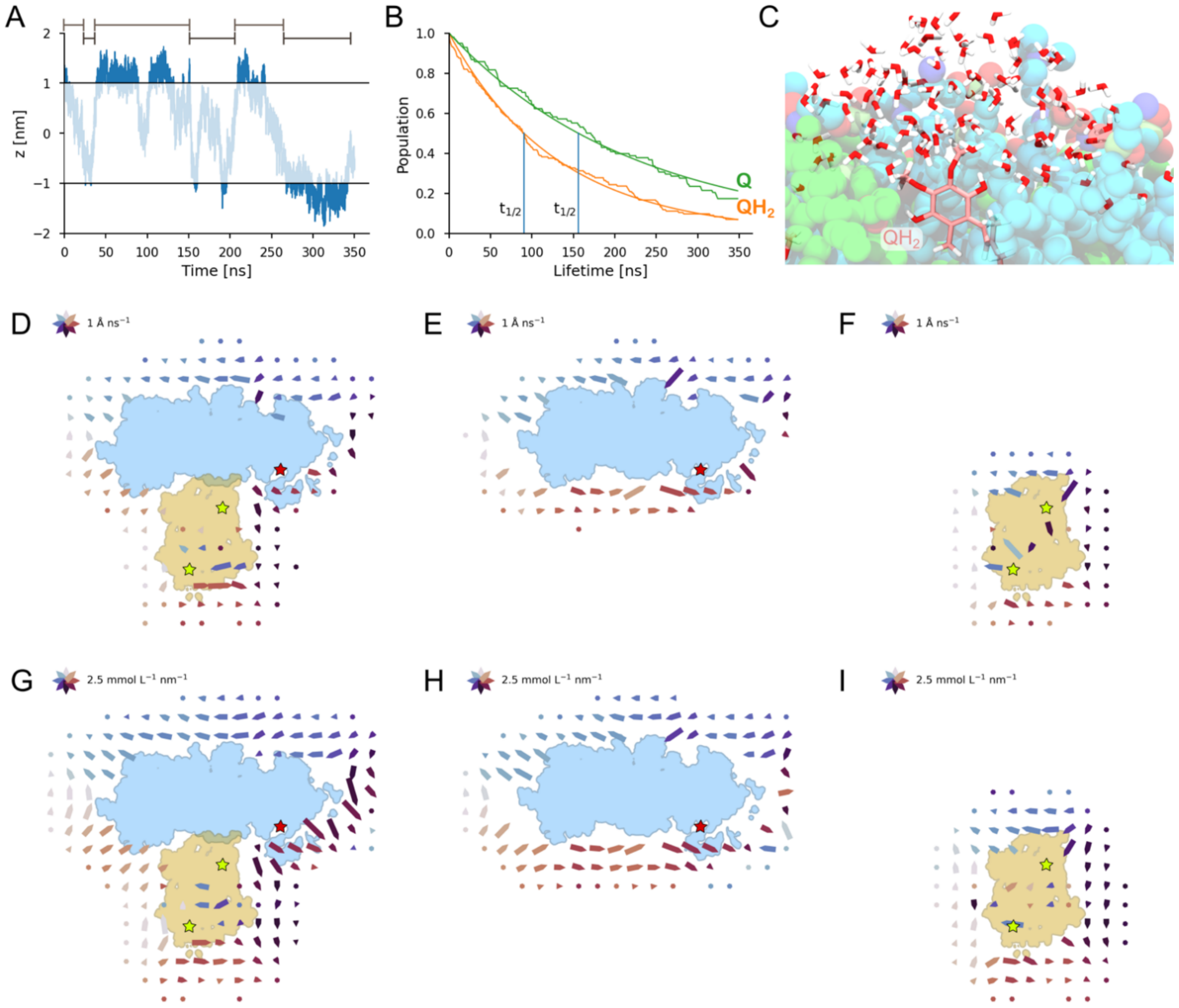


**Fig. S3.** Quinone dynamics in the membrane. (**A**) Flip-flop motion of the Q headgroup in the membrane from aMD simulations. Lifetime spent in upper/lower leaflets indicated by green bars. (**B**) Lifetime analysis of the Q and $QH_2$ headgroup, with the half-life ($t_{1/2}$) for Q/$QH_2$ spent on a given membrane leaflet during consecutive steps during the simulations. (**C**) Representative aMD structure of $QH_2$ headgroup interaction with water and lipid headgroups at the membrane interface. (**D**-**F**) Average diffusion of Q/$QH_2$ around the SC/IIII$_2$, CI, and CIII$_2$ from cgMD simulations. (**G**-**I**) Concentration gradient of Q/$QH_2$ around the SCI/III$_2$, CI, and CIII$_2$ from cgMD simulations. CI – in blue, CIII$_2$ – in beige. The Q species show a directed rotational diffusion around the OXPHOS complexes that could arise from conservation laws due to the Q depletion/increase in unique regions.

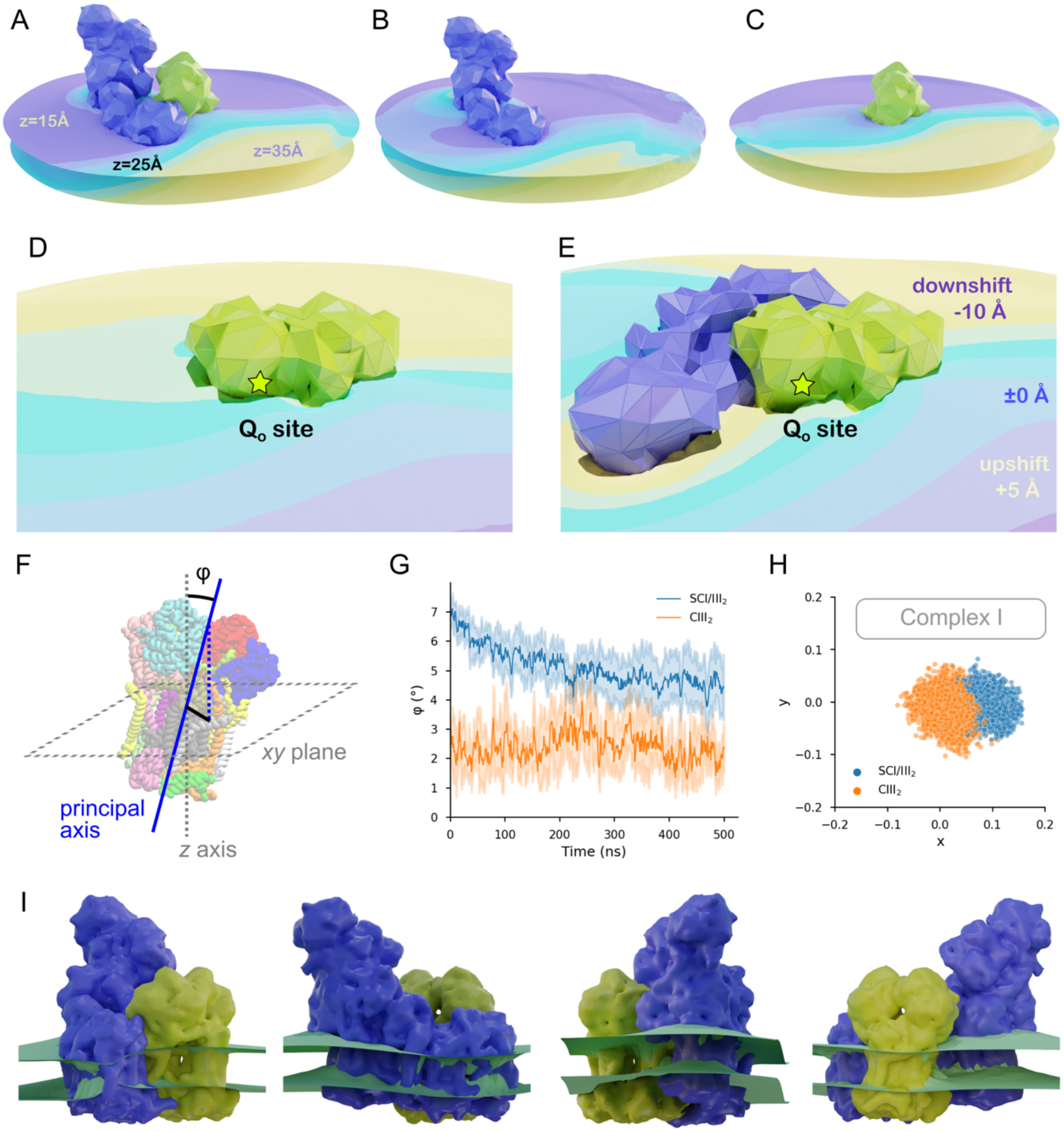


**Fig. S4.** Shifts in the membrane leaflet from cgMD simulations. (**A**-**C**) Shifts in the membrane leaflet viewed from the N-side of the (**A**) SCI/III$_2$, (**B**) CI, and (**C**) CIII$_2$. (**D**-**E**) Shifts in the membrane leaflet viewed from the P-side of the membrane for (**D**) CIII$_2$ and (**E**) SCI/III$_2$. The location of the Q$_o$ site of the proximal CIII protomer is indicated by a red star. (**F**) The CIII$_2$ angle was measured between the *z*-axis of the simulation box and the principal axis of inertia of the protein. (**G**) Time-evolution of the membrane angle averaged over aMD simulations of the SCI/III$_2$ and CIII$_2$. (**H**) Projection of the principal axis onto the *xy*-plane. (**I**) Visualization of the membrane distortion effect based on trajectory-averaged membrane position.

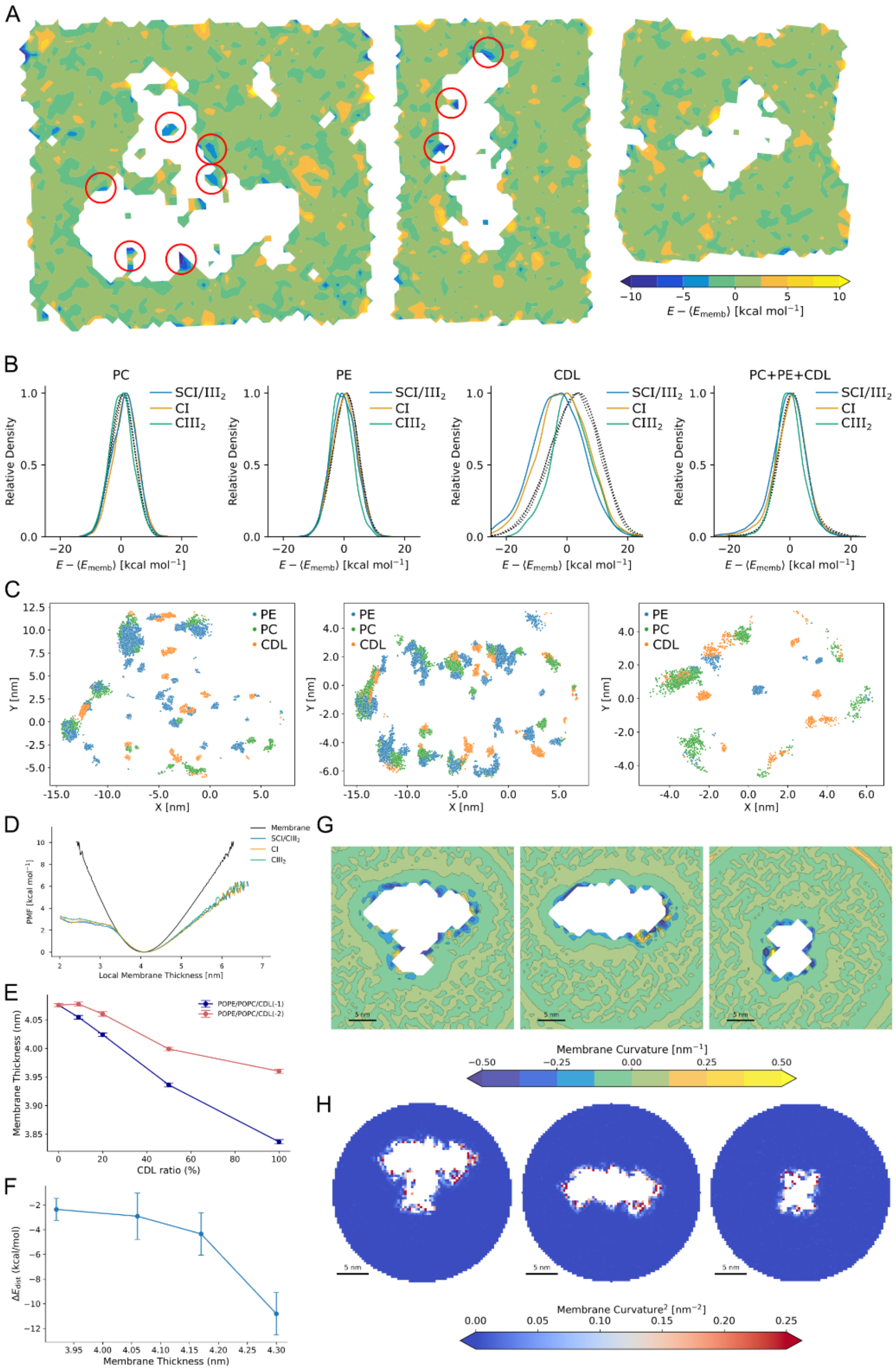


**Fig. S5.** Analysis of membrane-induced distortion effects. (**A**) Relative strain effect relative to a lipid membrane from atomistic MD simulations of the SCI/III$_2$, CI, and CIII$_2$, suggesting reduction of the membrane strain (blue patches) in the SC surroundings. The figure shows the non-bonded energies relative to the average non-bonded energies from membrane simulations (simulation M4, Table S1). (**B**) The lipid strain contribution for different lipids calculated from non-bonded interaction energies of the lipids relative to the average lipid interaction in a IMM membrane model (simulation M4). The figure shows the relative strain contribution for nearby lipids ($r$ < 2 Å, in color from panel (**C**), and lipids >5 Å from the OXPHOS proteins. (**C**) Selection of lipids (< 2 Å) interacting with the OXPHOS proteins. (**D**) Potential of mean force (PMF) of membrane thickness derived from thickness distributions from cgMD simulations of a membrane, the SCI/III$_2$, CI, and CIII$_2$. (**E**)

Membrane thickness as a function of CDL concentration from cgMD simulations. (**F**) $\Delta G_{thick}$ of the SC as a function of membrane thickness based on cgMD simulations. (**G**) Membrane curvature around the SCI/$III_2$ (*left*), CI (*middle*), and $CIII_2$ (*right*) from atomistic simulations. (**H**) Squared membrane curvature obtained from cgMD simulations, within a 20 nm radius around the center of the system. These maps correspond to the curvature field used in the calculation of the bending deformation energy term ($G_{curv}$).

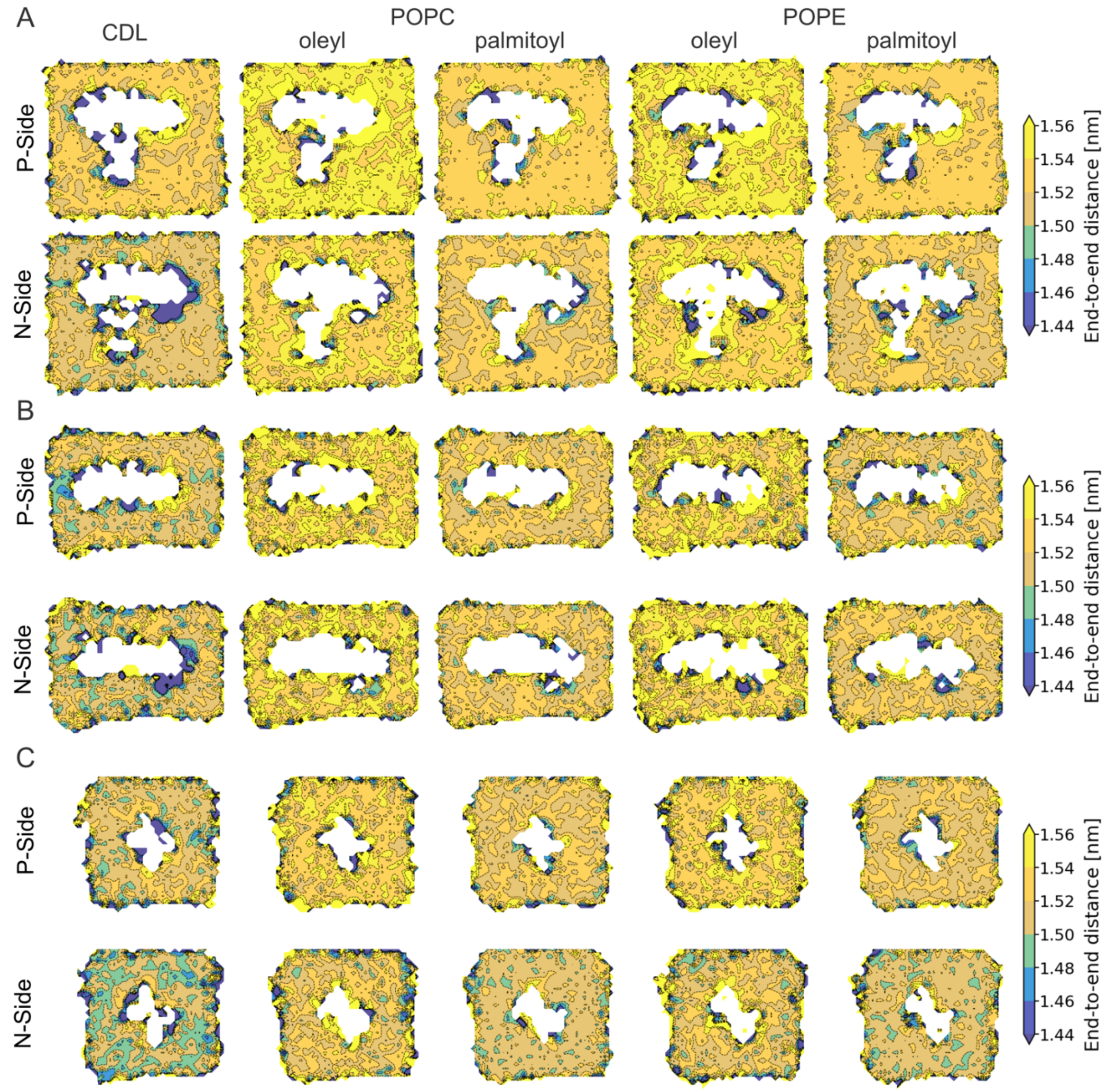


**Fig. S6.** Analysis of lipid *end-to-end* distance from aMD simulations of (**A**) SC, (**B**) CI, (**C**) $CIII_2$.

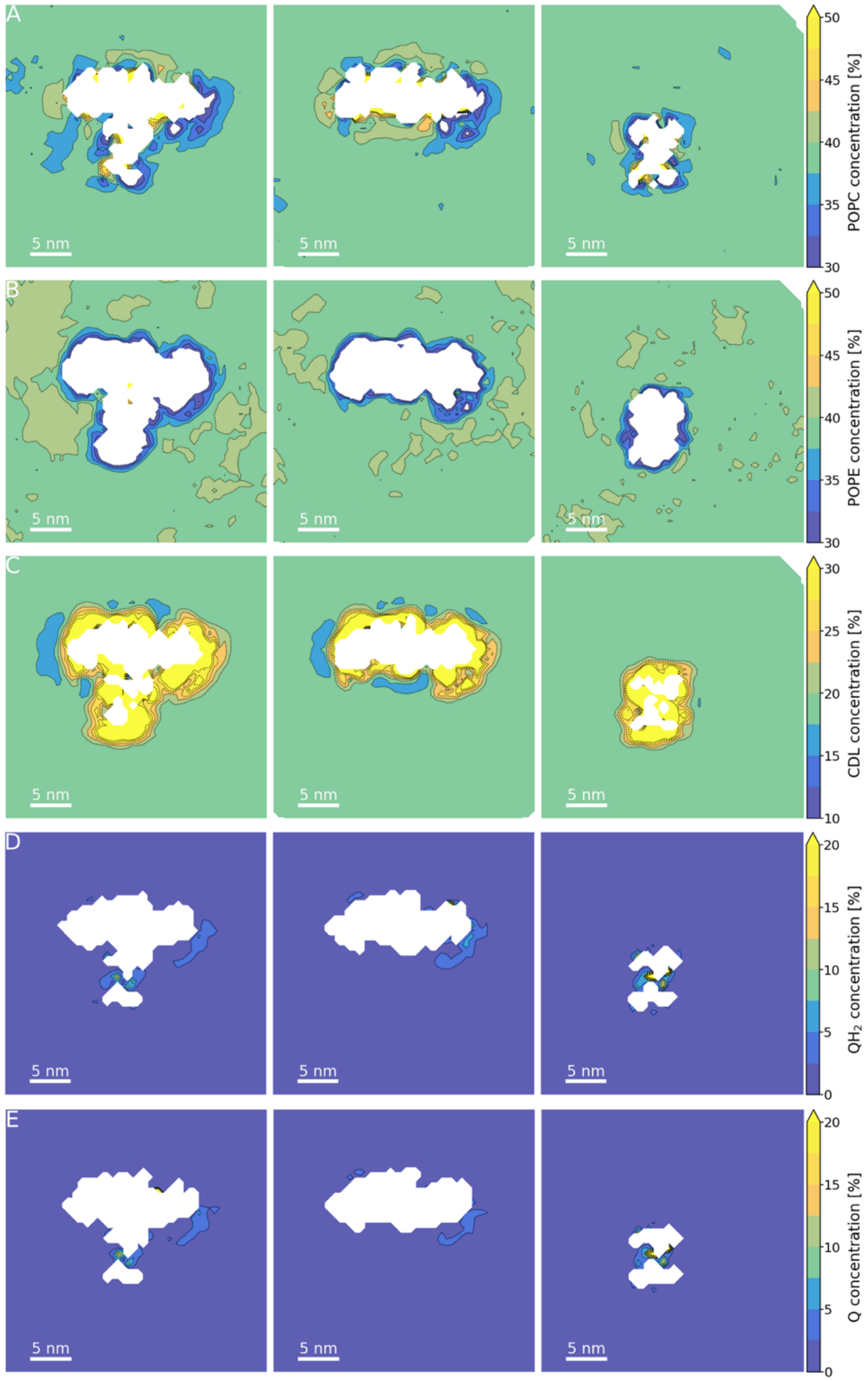


**Fig. S7.** Local lipid concentrations from cgMD simulations. Concentration of (**A**) POPC, (**B**) POPE, (**C**) CDL, (**D**) $QH_2$, and (**E**) Q (**E**) from cgMD simulations of the SCI/$III_2$ (*left*), CI (*middle*), and CIII (*right*).

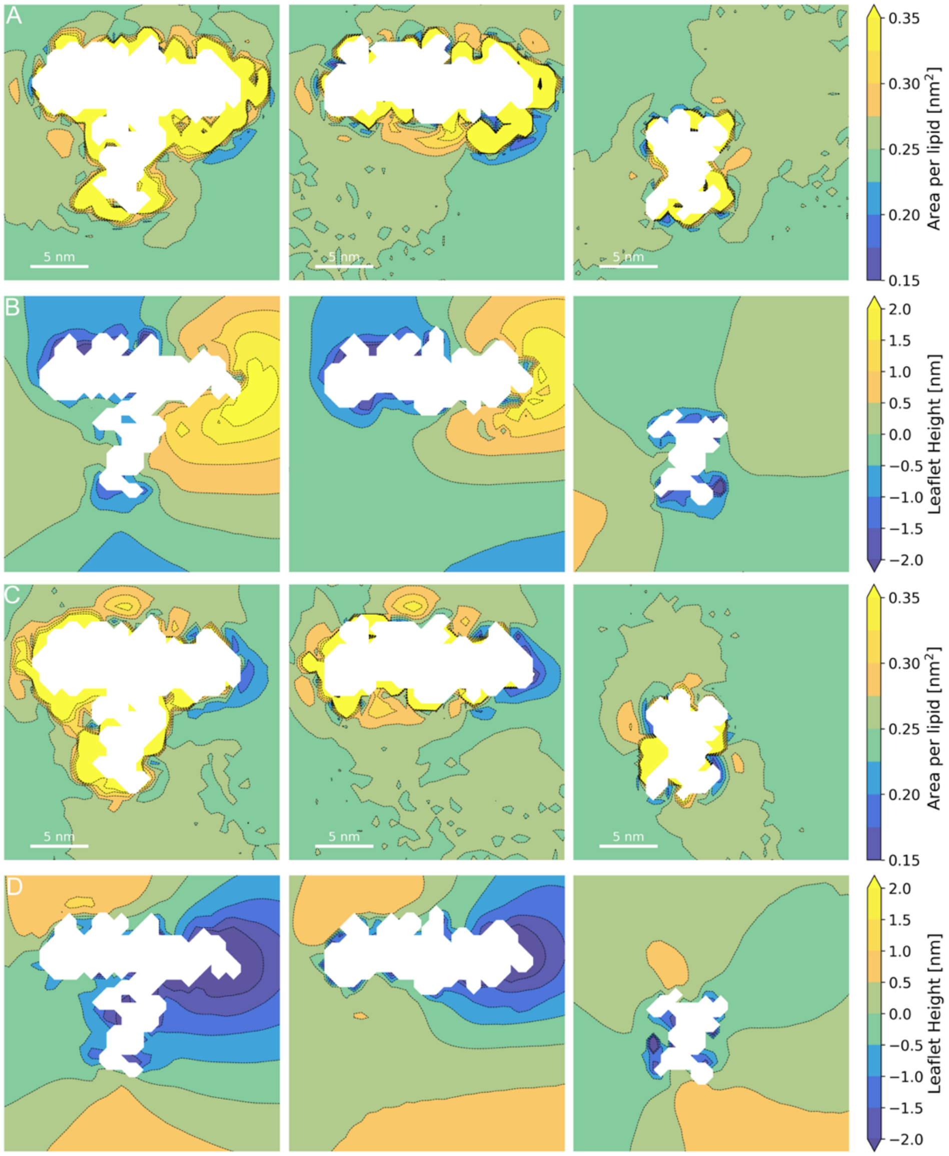


**Fig. S8.** Surface area per lipid and local distortion of membrane leaflet. (**A**) The area per lipid on the P-side of the membrane. (**B**) The local leaflet shift in the P-side of the membrane. (**C**) The area per lipid on the N-side of the membrane. (**D**) The local leaflet shift in the N-side of the membrane.

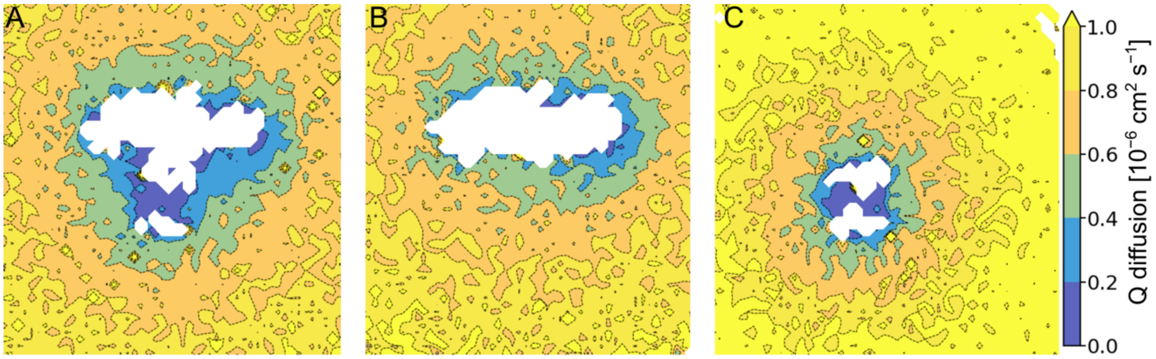


**Fig. S9.** Diffusion of Q molecules in cgMD simulations. (**A**-**C**) Map of Q diffusion for (**A**) SCI/III$_2$, (**B**) CI, and (**C**) CIII$_2$.

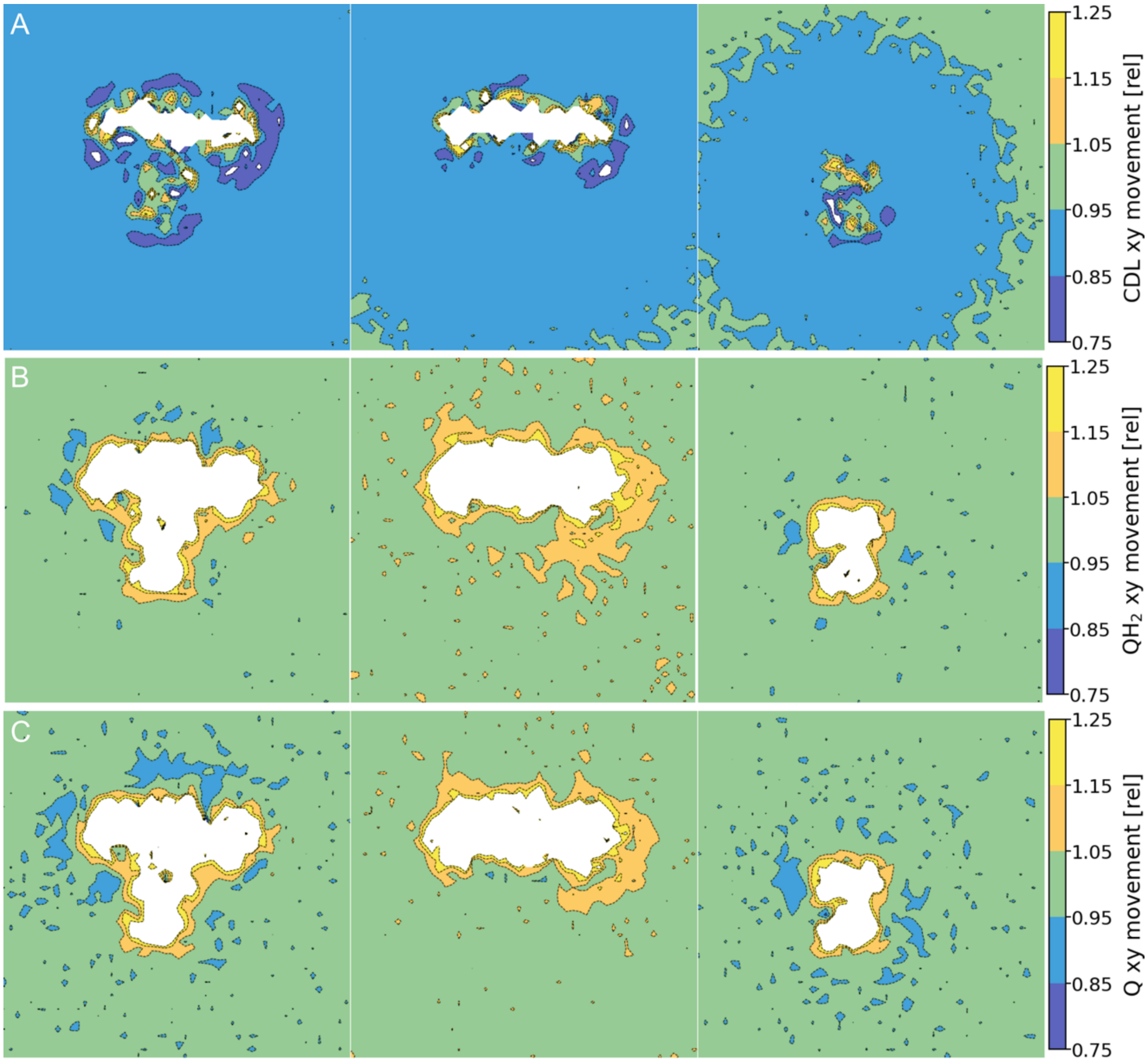


**Fig. S10.** Relative movement of lipids in cgMD simulations. (**A**-**D**) Map of average movement of (**A**) CDL and (**B**) $QH_2$ for SCI/$III_2$ (*left*), CI (*middle*), and $CIII_2$ (*right*). The movement is normalized relative to the local movement of POPC and POPE.

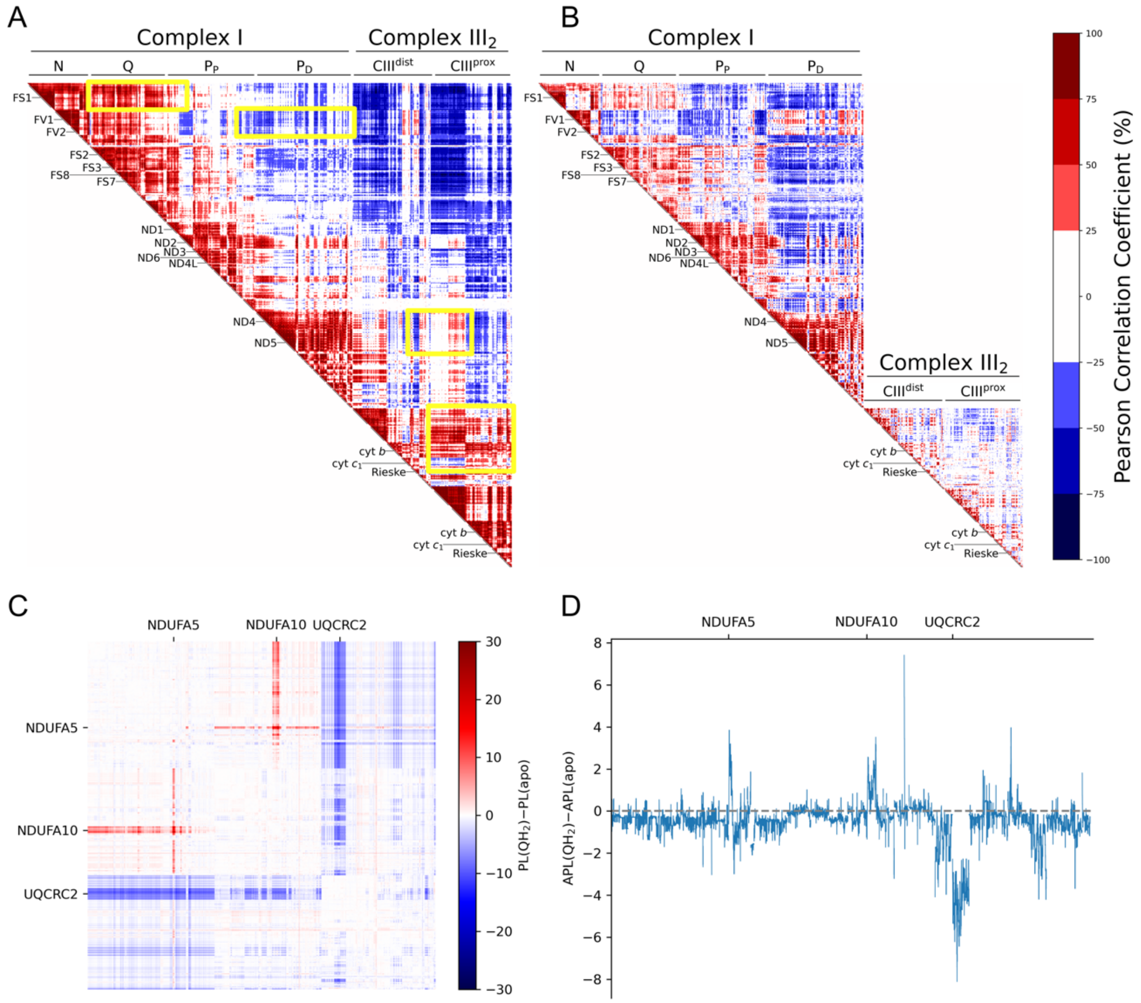


**Fig. S11.** Correlated motion within the SC and allosteric network analysis. (**A,B**) The Pearson correlation coefficients from Cα positions during aMD simulations for (**A**) within the SCI/III$_2$. (**B**) Correlation within the individual CI and CIII$_2$. Core subunits are indicated by labels. Differences between A and B are indicated by yellow boxes. (**C,D**) Allosteric network analysis (see *Supporting Methods*). (**C**) Path length difference of SCI/III$_2$ network for the CI QH$_2$ bound state and CI *apo* state. (**D**) Difference in average path length between the CI QH$_2$ bound state and CI *apo* state.

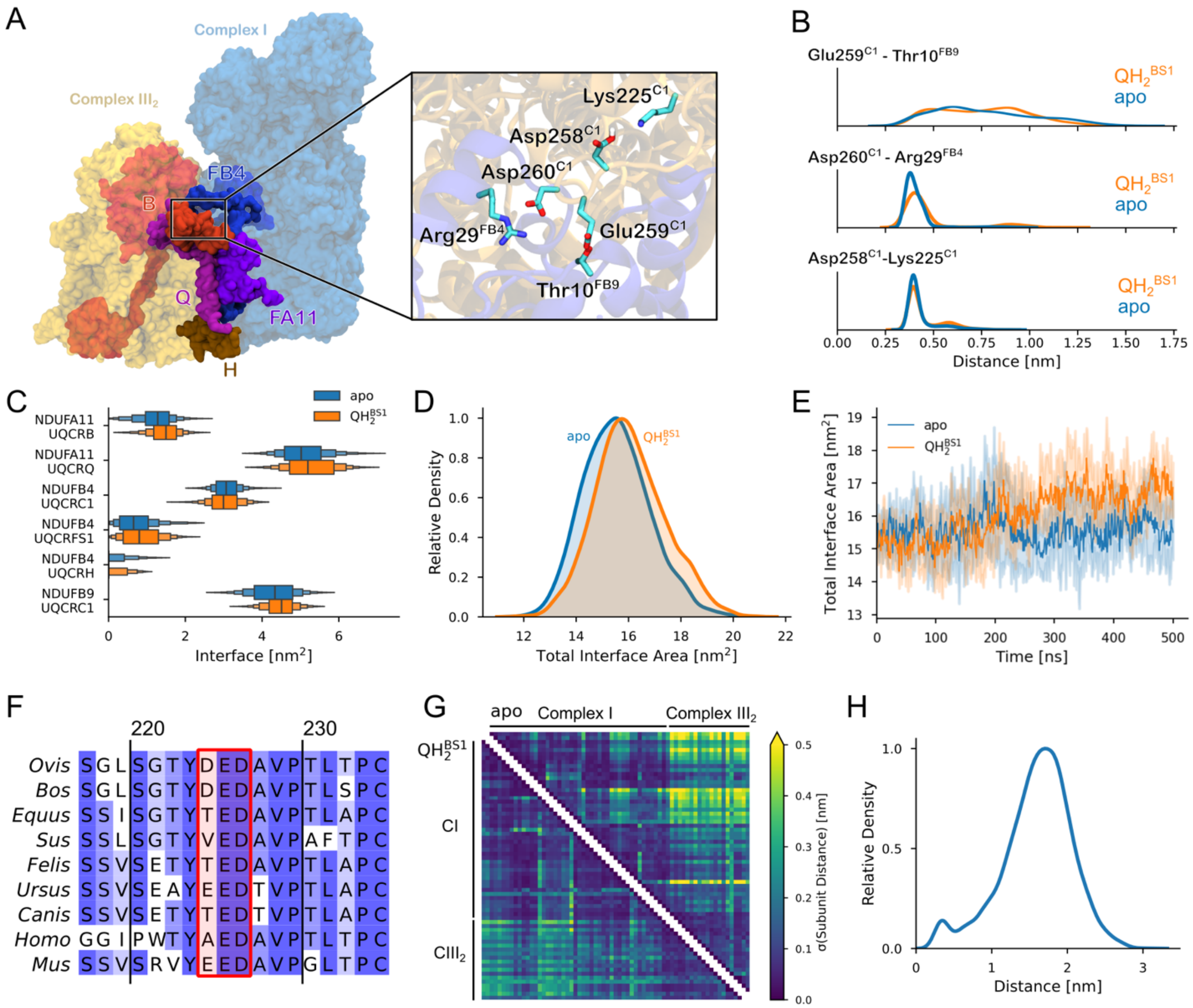


**Fig. S12.** Interaction area between CI and $CIII_2$ from aMD simulations. (**A**) Structure of the $SCI/III_2$ interface. *Inset:* Specific interactions at the CI and $CIII_2$ interface. (**B**) Distribution of the pair distances within the $SCI/III_2$. (**C**) Distribution of interface area between different subunits. (**D**) Distribution of the total interface area for simulations in the *apo* and $QH_2$-bound states of CI. (**E**) Total interface area as a function of simulation time. (**F**) Multiple sequence alignment of UQCRC1, with the carboxylate motif forming the SC contact highlighted in red. (**G**) Standard deviation of the inter-subunit distance matrix for the CI *apo* state (upper triangle) and CI $QH_2$ bound state (lower triangle). (**H**) Distance distribution between C-terminus of NDUFB7 and $CIII_2$. No specific contacts form between these regions during the MD simulations, supporting that substitution of this region resulted in unaltered SCs (12).

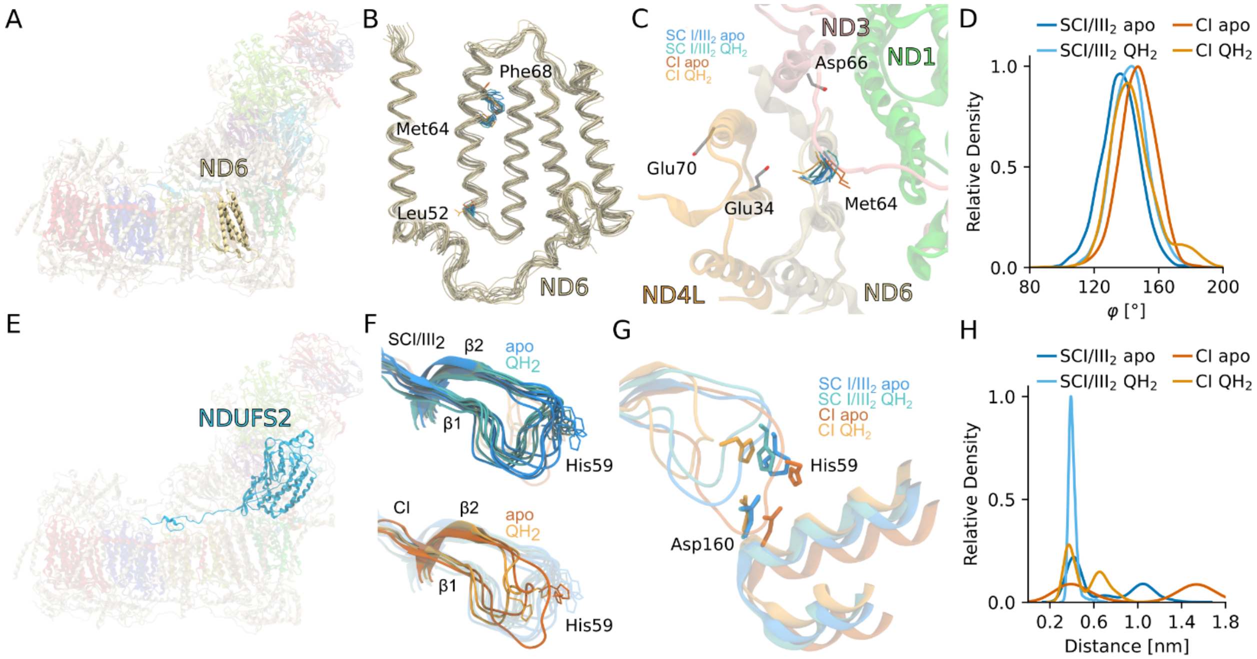


**Fig. S13.** Analysis of the $TM3^{ND6}$ dihedral angle in CI. (**A**) Position of the ND6 subunit within CI. (**B**,**C**) Average conformation of ND6 from last 100 ns - (**B**) the side view and (**C**) the top view of ND6. (**D**) Distribution of φ for $SCI/III_2$ and CI in the *apo* and $QH_2$-bound states of CI. (**E**) Position of the NDUFS2 subunit of CI. (**F**) Average conformation (over last 100 ns) of the β1-β2 loop of NDUFS2 from MD simulations for $SCI/III_2$ (*top*) and CI (*bottom*). (**G**,**H**) Distance distribution between His59 and Asp160 of NDUFS2.

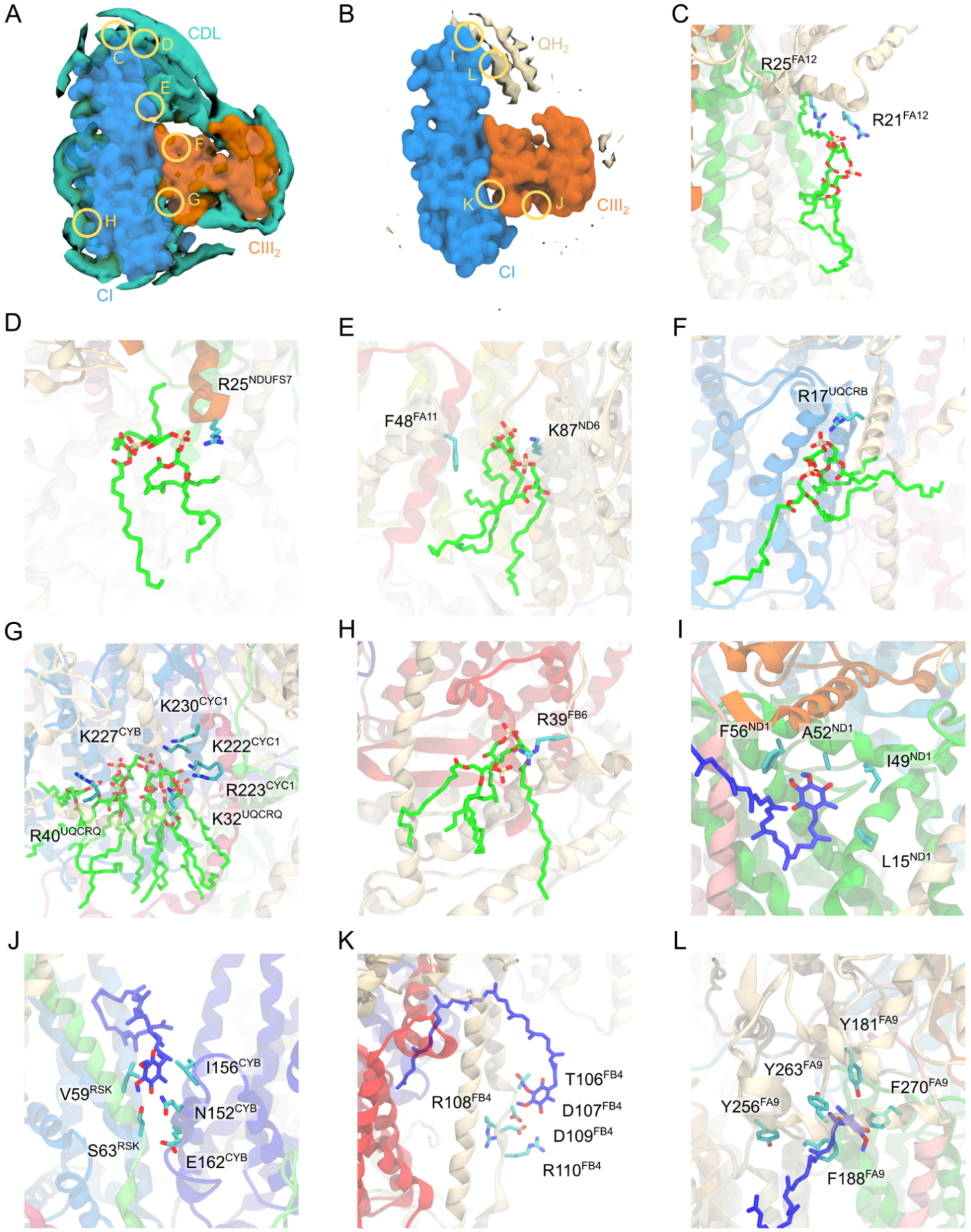


**Fig. S14.** Interaction of cardiolipin and quinone/quinol with the SCI/III2. (**A**) Overview over the CDL interaction sites show in (**C**-**H**). (**B**) Overview over the Q/QH2 interaction sites shown in (**I**-**L**).

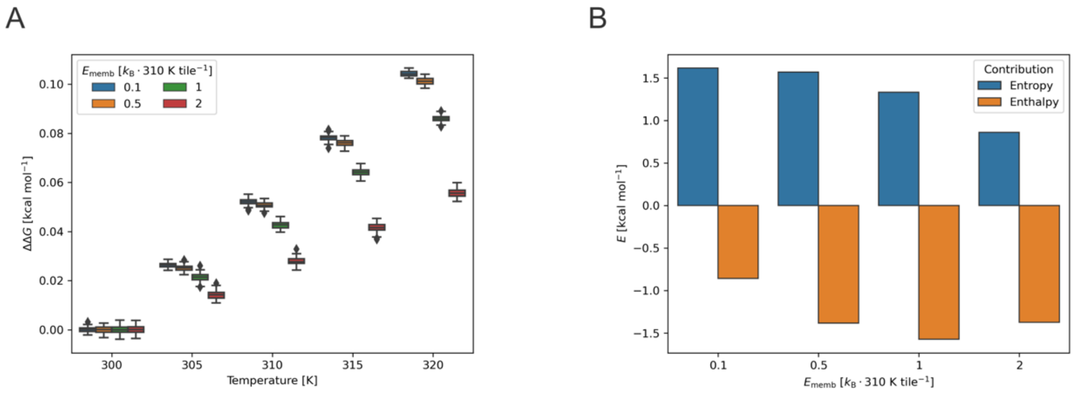


**Fig. S15.** Enthalpy-entropy compensations from the lattice model. (**A**) Free energy changes as a function of the introduced membrane strain term. (**B**) Enthalpic and entropic contributions to the free energy as a function of the introduced membrane strain term.

**Table S1.** List of atomistic MD and cgMD simulations.

**aMD:**

| **Simulation Name** | **Protein Model** | **CI Ligand State** | **CIII$_{proximal}$ Ligand State** | **CIII$_{distal}$ Ligand State** | **Simulation Time (μs)** |
|---|---|---|---|---|---|
| A1 | SCI/III$_2$ | *apo* | *apo* | *apo* | 2 × 0.5 |
| A2 | | *apo* | Q in $Q_i$, $QH_2$ in $Q_o$ | *apo* | 2 × 0.5 |
| A3 | | *apo* | *apo* | Q in $Q_i$, $QH_2$ in $Q_o$ | 2 × 0.5 |
| A4 | | $QH_2$ | *apo* | *apo* | 2 × 0.5 |
| A5 | | $QH_2$ | Q in $Q_i$, $QH_2$ in $Q_o$ | *apo* | 2 × 0.5 |
| A6 | | $QH_2$ | *apo* | Q in $Q_i$, $QH_2$ in $Q_o$ | 2 × 0.5 |
| A7 | CI | $QH_2$ | *n/a* | *n/a* | 2 × 0.5 |
| A8 | CI | *apo* | *n/a* | *n/a* | 2 × 0.5 |
| A9 | CIII$_2$ | *n/a* | *apo* | *apo* | 2 × 0.5 |
| A10 | | *n/a* | Q in $Q_i$, $QH_2$ in $Q_o$ | *apo* | 2 × 0.5 |
| A11 | | *n/a* | *apo* | Q in $Q_i$, $QH_2$ in $Q_o$ | 2 × 0.5 |
| | | | | **Total:** | **11 μs** |

**Table S1 (contd.).** List of atomistic MD and cgMD simulations.

**cgMD:**

| Simulation Name | Protein Model | CI Ligand State | $CIII_{proximal}$ Ligand State | $CIII_{distal}$ Ligand State | Simulation Time (μs) |
|---|---|---|---|---|---|
| C1 | $SCI/III_2$ | $QH_2$ | Q in $Q_i$, $QH_2$ in $Q_o$ | Q in $Q_i$, $QH_2$ in $Q_o$ | 75 + 50 |
| C2 | CI | $QH_2$ | *n/a* | *n/a* | 50 + 40 |
| C3 | $CIII_2$ | *n/a* | Q in $Q_i$, $QH_2$ in $Q_o$ | Q in $Q_i$, $QH_2$ in $Q_o$ | 50 + 50 |
| | | | | **Total:** | **315 μs** |

**Table S2.** Simulation details of membrane models.

**aMD:**

| Simulation Name | System size (Å$^3$) | Lipid Composition | Lipid/Q/QH$_2$ Molecules | Simulation Time (μs) |
|---|---|---|---|---|
| M1 | 78×78×82 | POPC/POPE (1:1) | 204 | 2 × 0.2 |
| M2 | 78×78×82 | CDL | 102 | 2 × 0.2 |
| M3 | 187×187×182 | POPC/POPE/CDL/Q (38:38:19:5) | 1000 | 2 × 0.35 |
| M4 | 187×187×182 | POPC/POPE/CDL/QH$_2$ (38:38:19:5) | 1000 | 2 × 0.35 |

**cgMD:**

| Simulation Name | System | Lipid Bead Distance (nm) | Simulation Time (μs) |
|---|---|---|---|
| cgM1/2/3 | SCI/III$_2$ | 0.44 / 0.50 / 0.53 | 5 / 5 / 5 |
| cgM4/5/6 | CI | 0.44 / 0.50 / 0.53 | 5 / 5 / 5 |
| cgM7/8/9 | CIII$_2$ | 0.44 / 0.50 / 0.53 | 5 / 5 / 5 |
| cgM10-13 | Membrane | 0.44 / 0.47 / 0.50 / 0.53 | 5 / 23 / 5 / 5 |

**Table S3.** Non-standard protonation states used in aMD simulations. *The protonation states of $H59^{NDUFS2}$ and $Y108^{NDUFS2}$ were modeled in their neutral ($His^0$)/deprotonated ($TyrO^-$) states with $QH_2$ in CI.

| Subunit | Residues |
|---|---|
| ND1 | E192, E206, H247(ε), H287(ε) |
| ND2 | H25(ε), K46, H48(ε), H112(ε), H186(ε), H232(ε), K263 |
| ND4 | H82(ε), H213(ε/δ), H220(ε), Lys283, H293(ε), H319(ε), H338(ε/δ), H419(ε), H422(ε) |
| ND5 | H27(ε), H56(ε), H109(ε), K119, H230(ε), H248(ε), H323(ε), H348(ε), K392, H484(ε), H509(ε), H605(ε) |
| NDUFS2 | H55(ε), H59(ε/δ)*, Y108*, D104, H150(ε), H157(ε), H190(ε), H200(ε), D292, E343, H348(ε) |
| ND3 | D66, E68, E105 |
| NDUFS7 | D68, E154 |
| NDUFS3 | H19(ε), H53(ε), H145(ε) |
| NDUFV2 | H9(ε), H42(ε), H99(ε) |
| NDUFV1 | H29(ε), D98, H113(ε), H116(ε), H261(ε/δ), H283(ε), H356(ε/δ), H437(ε) |
| NDUFS1 | H43(ε), D232, H255(ε), H293(ε/δ), D324, E347, H401(ε), H421(ε), H437(ε), H494(ε), H549(ε) |
| NDUFS8 | H65(ε/δ), K81, H144(ε/δ) |
| ND6 | E100 |
| ND4L | H52(ε/δ) |
| 18 kDa | H29(ε) |
| 9 kDa | H43(ε), H44(ε), H75(ε) |
| B8 | H21(ε) |
| B12 | Glu29 |
| B17 | H67(ε), H74(ε), H83(ε), H89(ε), H127 (ε/δ) |
| B18 | H3(ε), H60(ε), HSP81, H84(ε), Asp87, Glu90, H91(ε) |
| B22 | H11(ε), H25(ε), H32(ε), H50(ε), H72(ε), H75(ε), H107(ε) |
| AGGG | H6(ε), H42 (ε/δ), H50(ε) |
| ASHI | H66(ε), H78(ε), H104(ε), H155(ε) |
| ESSS | H45(ε) |
| MNLL | H10(ε), H13(ε) |

**Table S4.** Estimation of volume changes from cgMD simulations. The volume change of the membrane with embedded OXPHOS proteins. The SCI/$III_2$ formation leads to the membrane strain relative to the individual CI and $CIII_2$.

| **System** | **Membrane volume ($nm^3$)** | **Δ*V* ($nm^3$)** |
|---|---|---|
| Membrane | 4160 ± 1.0 | |
| CI | 3946 ± 0.8 | 214 |
| $CIII_2$ | 4093 ± 0.7 | 68 |
| SCI/$III_2$ | 3865 ± 0.7 | 296 |
| **Δ*V*:** | | **14 $nm^3$** |

**Table S5.** Estimation of protein copy numbers and effective surface area in the IMM. The data is based on Refs. 7, see also Refs. 86-90. The crowding model, with a square length of 163 nm, describes the flat membrane regions of the IMM, thus excluding the area of ATP synthase and 10 lipids bound the c-ring, located at the cristae (*cf.* Ref. 86).

| **Component** | **Copy number** | **Surface area/molecule ($nm^2$)** | **Total surface area ($nm^2$)** |
|---|---|---|---|
| CI | 13[86] | 190 | 2470 |
| CII | 15[87] | 16 | 240 |
| $CIII_2$ | 18[86] | 110 | 1980 |
| CIV | 64 | 64 | 4096 |
| ATP synthase | 21[88] | 200 | 4200 |
| ATP/ADP carrier | 160[89] | 10 | 1600 |
| Lipids | 48,300[90] | 0.7 | 16,905/leaflet |
| of which $Q/QH_2$ 1% | 483 | | |
| | **291 proteins 48,300 membrane** | | **14,586 $nm^2$ protein 16,905 $nm^2$ membrane Tot: 31,491 $nm^2$** |

**Table S6**. Energy terms in the lattice model. The protein-protein interaction is described by specific interactions term ($E_{specific} < 0\ k_BT$) and non-specific interactions ($E_{non\text{-}specific}>0$). The membrane-protein interaction determines the strain energy of the membrane ($E_{strain}$), based on the number of neighboring "lipid"-occupied grids that are in contact with proteins (Fig. 4A). The interaction between the lipids was indirectly accounted for by the background energy of the model. The proteins can occupy four unique orientations on a grid ([North, East, South, West]). The table summarizes the unique energies linked to the respective microstates.

| State | CI and $CIII_2$ position | $d(CI - CIII_2)$ | Energy |
|---|---|---|---|
| 1 | Neighbors with specific interaction | 1 | $E_{specific} + 10\ E_{strain}$ |
| 2 | Neighbors with non-specific interaction | 1 | $E_{non\text{-}specific} + 10\ E_{strain}$ |
| 3 | Diagonal neighbors | $\sqrt{2}$ | $12\ E_{strain}$ |
| 4 | Separated by one lattice position in either cardinal direction | 2 | $13\ E_{strain}$ |
| 5 | Separated by one lattice position in cardinal direction | $\sqrt{5}$ | $14\ E_{strain}$ |
| 6 | Diagonal neighbors separated by one lattice position | $2\sqrt{2}$ | $15\ E_{strain}$ |
| 7 | No interaction | $> 2\sqrt{2}$ | $16\ E_{strain}$ |

**Table S7.** Monte Carlo simulations of the lattice model. The conformational landscape was sampled by Monte Carlo (MC) using $10^7$ MC iterations with 100 replicas. Temperature effects were modeled by varying $\beta$, and the effect of different *protein*-to-*lipid* ratios by increasing the grid area. The following simulations were performed (energy units are given in $k_B \times 310$ K):

| Simulation | Temperature (K) | Grid size $N$ | $E_{specific}$ | $E_{non\text{-}specific}$ | $E_{strain}$ |
|---|---|---|---|---|---|
| G1-G5 | 300, 305, …, 320 | 4 | -1 | 1 | 0.1 |
| G6-G10 | 300, 305, …, 320 | 4 | -1 | 1 | 0.5 |
| G11-G15 | 300, 305, …, 320 | 4 | -1 | 1 | 1 |
| G16-G20 | 300, 305, …, 320 | 4 | -1 | 1 | 2 |

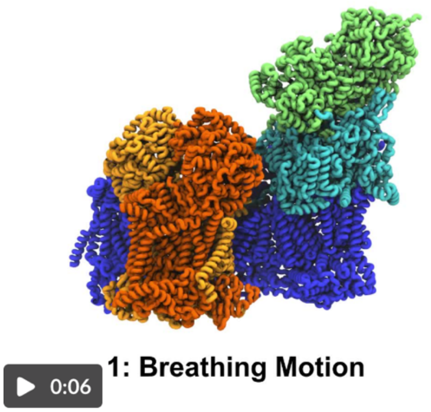


**Movie S1.** Normal modes of the SC.

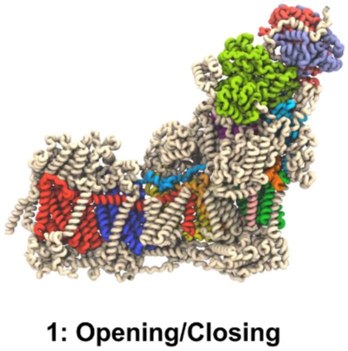


**Movie S2.** Normal modes of the isolated CI.

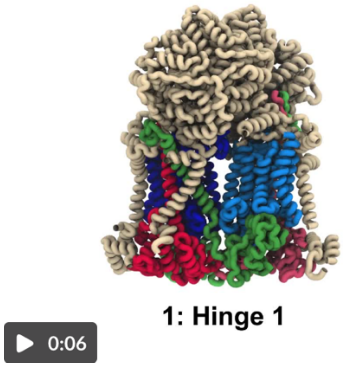


**Movie S3.** Normal modes of the isolated $CIII_2$.

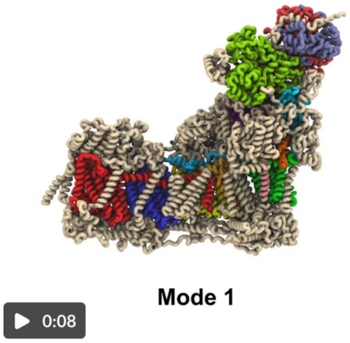


**Movie S4.** Comparison of the normal modes between the SC and the isolated CI.

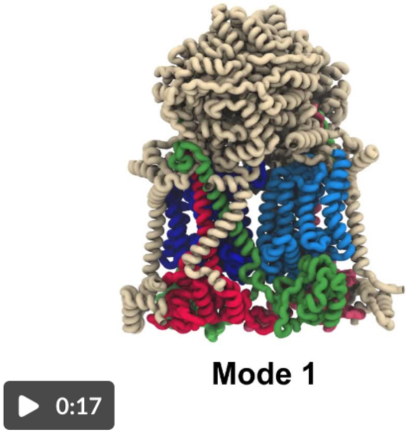


**Movie S5.** Comparison of the normal modes between the SC and the isolated $CIII_2$.
